# In-depth study of MPV17: a molecular travel unveiling a mitochondrial calcium regulation function

**DOI:** 10.1101/2024.04.12.589178

**Authors:** Sébastien Meurant, Ilario Amato, Lorris Mauclet, Marc Dieu, Arnaud Chevrollier, Benjamin Ledoux, Marino Caruso, Guy Lenaers, Thierry Arnould, Patricia Renard

**Affiliations:** URBC, Namur Research Institute for Life Sciences (Narilis), University of Namur (UNamur), 5000, Namur, Belgium; Mass Spectrometry Platform (MaSUN), University of Namur (UNamur), 5000, Namur, Belgium; Univ Angers, CNRS UMR 6015, INSERM U1083, SFR ICAT, Angers, France; SCiAM platform, SFR ICAT, Université d’Angers, Angers, France; Morphology & Imaging Platform (Morph-Im) - Namur Research Institute for Life Sciences (Narilis), University of Namur (UNamur), 5000, Namur, Belgium; Laboratory for Disease Mechanisms in Cancer, Department of Oncology, KU Leuven, Leuven, Belgium; Service de Neurologie, CHU d’Angers, Angers, France

**Keywords:** BioID, Calcium regulation, Mitochondria, MICOS, mPTP, MPV17, mtDNA maintenance

## Abstract

Mitochondrial DNA depletion syndromes are severe genetic disorders associated with mutations in a variety of genes including *MPV17,* encoding a protein of the inner mitochondrial membrane with an unclear function. In this study, using BioID technology, we identified MPV17 interacting partners among which proteins from the MICOS complex. However, MPV17 knockout did not impact mitochondrial ultrastructure, but led to increased mitochondria-derived vesicles formation and altered mitochondrial permeability transition pore. Furthermore, MPV17 KO cells exhibited higher mitochondrial calcium levels and reactive oxygen species, leading to mtDNA degradation, a phenomenon prevented by blocking mitochondrial calcium entry or treating cells with antioxidant. We thus propose a function for MPV17 as a potential additional member of the mitochondrial permeability transition pore, whereas in the absence of the protein, the build-up of calcium inside the mitochondria would lead to mtDNA degradation caused by increased oxidative damages.

## 1. INTRODUCTION

The mitochondrial protein MPV17 is a five transmembrane domains-containing protein spanning the inner mitochondrial membrane. Mutations in this protein are involved in rare severe mitochondrial diseases, namely mitochondrial DNA (mtDNA) depletion syndromes (MDDS), also called mitochondrial DNA maintenance defect (MDMD), and Navajo neuro-hepatopathy (El-Hattab et al., 2018; Spinazzola et al., 2006). In addition, *MPV17* mutations are also described in Charcot-Marie-Tooth disease and axonal neuropathy (Blakely et al., 2012; Choi et al., 2015; Zaman et al., 2023). Genes associated with MDDS/MDMD are involved in different mitochondrial processes such as the mtDNA replication (*POLG*, *POLG2*, *TWK*, *TFAM*, *RNASEH1*, *MGME1* and *DNA2*), the maintenance and supply of the mitochondrial nucleotide pool (*TK2*, *DGUOK*, *SUCLG1*, *SUCLA2*, *ABAT*, *RRM2B*, *TYMP*, *SLC25A4* and *AGK*) and the mitochondrial fusion (*OPA1*, *MFN2* and *FBXL4*) (El-Hattab et al., 2017; Suomalainen & Isohanni, 2010). MPV17 is thus associated with mtDNA maintenance, but its molecular function is still unclear. The protein was proposed to be involved in the purine mitochondrial salvage pathway of main importance in quiescent cells to ensure the maintenance of the mitochondrial deoxynucleotide pool (Dalla Rosa et al., 2016). Additionally, expression of MPV17 mutants in HEK293T cells decreases mitochondrial bioenergetics (Jacinto et al., 2021). Besides, results from experiments inserting the human MPV17 protein into lipid membranes disclosed a non-selective channel function with an affinity for cations, and a role in maintaining the membrane potential under stress conditions (Antonenkov et al., 2015). However, another study revealed that MPV17 acts as a channel modulating the selective entry of metabolites such as uridine or orotate, and is sensitive to oxidative stress (Corra et al., 2023; Mukherjee et al., 2023; Sperl & Hagn, 2021). Upon oxidative stress, the protein could oligomerize, promoting the selective entry of metabolites for mtDNA repair or replication (Sperl & Hagn, 2021).

SYM1, the yeast ortholog of MPV17, exhibits strong evolutionary conservation as its depletion can be complemented by human MPV17 expression which can rescue the impaired OXPHOS activity and growth defect under non-fermentable condition (Dallabona et al., 2010). SYM1 depletion leads to mitochondrial ultrastructure defects (Dallabona et al., 2010), as also observed in glomeruli cells of *MPV17^-/-^*-related kidney disease mouse model (Luna-Sanchez et al., 2020) and in *mpv17^-/-^* zebrafish larvae muscle (Bian et al., 2021).

The maintenance of the mitochondrial ultrastructure is mainly achieved by the MItochondrial contact site and Cristae Organizing System (MICOS) complex, which is part of the Mitochondrial Intermembrane space Bridging (MIB) supercomplex. The MICOS complex is involved in the cristae formation and cristae junction stabilization and bridges the outer and inner membranes of mitochondria, by the interaction with the SAM50 outer membrane complex (reviewed in Kozjak-Pavlovic, 2017). Interestingly, MICOS disassembly and destabilization of the mitochondrial ultrastructure were reported in glomeruli cells from *MPV17^-/-^*mouse model, and OPA1 overexpression restored the mtDNA depletion observed in those cells (Luna-Sanchez et al., 2020). Additionally, MPV17 co-immunoprecipitates with the MICOS complex as well as with components of the ATP synthase complex and the membrane Permeability Transition Pore (mPTP) in mouse cardiomyocytes. Functionally, MPV17 was also reported to be associated with the mitochondrial calcium (Ca^2+^) retention capacity and with the mPTP regulation under oxidative stresses in mouse heart ischemia-reperfusion condition (Madungwe et al., 2020). Importantly, mitochondrial Ca^2+^ homeostasis depends on the balanced cross-talk between its entry and exit out of the mitochondria and is mainly ensured by the Mitochondrial Ca^2+^ Uniporter (MCU) Multi-Protein Complex while its efflux is controlled by different transporters including the mPTP (reviewed in Bernardi et al., 2023; Tanwar et al., 2021). Increase in calcium level can lead to Reactive Oxygen Species (ROS) production (Brookes et al., 2004) and ROS can damage mtDNA which can be repaired by mitochondrial machinery or degraded either by resident nucleases (Cullen & Steinberg, 2018; Fontana & Gahlon, 2020; Fu et al., 2020; Zhao, 2019) or selectively removed out of mitochondria and targeted to the lysosome through Mitochondria-Derived Vesicles (MDVs) (Picca et al., 2020; Sugiura et al., 2014). The main mtDNA quality control mechanism triggering the selective removal of oxidative damaged mtDNA molecules involves, in mouse, the VPS35-SAM50-ATAD3-TWNK axis, promoting the formation of MDVs and their degradation in the lysosome thanks to the endosomal system (Sen et al., 2022). A similar degradation pathway for oxidized mtDNA was also described in human and involves the primate-specific ATAD3B mitophagy-receptor (Shu et al., 2021).

Finally, another interesting feature of the yeast ortholog of MPV17 which could bring additional light regarding its function in human is its association with a >600 kDa complex (Dallabona et al., 2010). Interestingly, several mutants of SYM1, reminiscent of human MPV17 pathogenic alleles, have been shown to dissociate from this complex in yeast (Gilberti et al., 2018). This phenotype is also retrieved in *MPV17^-/-^* mice wherein the liver complementation with WT human MPV17 restores liver mtDNA levels, and the association of MPV17 with the macromolecular complex is correlated with the rescue of the diet-induced liver failure observed in those mice (Bottani et al., 2014). The identification of the MPV17-containing complex might thus be of interest to decipher the function of the inner mitochondrial membrane protein.

In this study, we characterized the proteins located in the close vicinity of MPV17 of HEK293T cells using proximity labelling (BioID) method (Roux et al., 2012). Our results support an association of MPV17 with the MICOS complex, although no mitochondrial ultrastructure defects were observed in MPV17 KO cells. Additional *in silico* analyses and modeling of MPV17 highlighted an association of the protein with the mPTP regulator CypD, which was confirmed experimentally. Interestingly, higher mitochondrial calcium level is observed in KO cells which also display a higher calcium threshold for mPTP opening. This higher calcium concentration/calcium accumulation is associated with higher ROS levels, and both the blockade of calcium entry and antioxidant treatment can rescue the mtDNA depletion observed in MPV17 KO cells. We thus identified a new function for MPV17, a protein that, in interaction with the mPTP, regulates mitochondrial calcium homeostasis.

## 2. MATERIAL AND METHODS

### 2.1 Cell culture and treatment

The HEK293T cells (obtained from ATCC) were cultured in Dulbecco’s Modified Eagle Medium (DMEM) with 4.5 g/L glucose and supplemented with 10 % Fetal Bovine Serum (FBS) (ThermoFisher Scientific, Waltham, USA) and were kept at 37 °C and 5 % CO_2_. Expanding cells were maintained under 80 % confluency and passed by using trypsin-EDTA (ThermoFisher Scientific). For the autophagy inhibition, cells were treated with 50 nM bafilomycin A1 (Sigma-Aldrich, Saint-Louis, USA) or 5 µM MRT68921 HCl (Selleck Chemicals, Houston, USA) for 48 h. For MCU inhibition, cells were treated with 5 µM Ru360 (Selleck Chemicals) for 72 h. For antioxidant treatment, cells were treated with 4 mM N-Acetyl-L-Cysteine (NAC) (Sigma-Aldrich) for 72 h. For each treatment or transfection, in order to be able to observe the mtDNA depletion phenotype at confluency, the cells were treated for the last 24 h of treatment with 10 nM ethidium bromide (Sigma-Aldrich) supplemented with 50 μg/mL uridine (Sigma-Aldrich) and 1 mM pyruvate (ThermoFisher Scientific), to block mtDNA replication.

### 2.2 Generation of KO cell lines

KO of *MPV17* in HEK293T cells was performed using CRISPR-Cas9 technology as described previously (Meurant et al., 2023). Briefly, crRNAs targeting the exons 2 (5’-TTTCCACGGGTGAGCGGCCAGGG-3’) (Integrated DNA Technologies (IDT), Coralville, USA) or 4 (5’-GGTTTGGATCGGTTCATCCCTGG-3’) of the *MPV17* gene were designed using CRISPOR bioinformatic resource (Concordet & Haeussler, 2018) and used separately or in combination. 150 picomoles of crRNA were annealed 1:1 with a universal tracrRNA (IDT) by cooling down from 95 °C to 30 °C at a rate of 5 °C per min. The crRNA:tracrRNA heteroduplex was then mixed with 10 µg of recombinant Cas9 (IDT) for 10 min at room temperature. Meanwhile, 1 million cells were harvested and resuspended in nucleofector solution from Nucleofector Kit V (Lonza), supplemented with the ribonucleoparticle of crRNA:tracrRNA and Cas9. Cells were next electroporated using Amaxa Nucleofector Ib with the C-09 program and plated in 6-well plates with prewarmed medium. After 72 h, cells were passed with a limiting dilution and single cells were transferred in a 96-well plate. The clones were then expanded and three different WT and four different KO clones were selected based on Sanger sequencing screening (Eurofins Discovery, St. Charles, USA). Additionally, the KO were confirmed following the assessment of MPV17 abundance by mass spectrometry (**Table EV1**).

### 2.3 Vectors cloning

The BioID vectors, MPV17-BirA*, preOTC-BirA* and TWNK-BirA*, were constructed using gateway cloning, as described previously (Liu et al., 2018). Briefly, the Coding DNA Sequence (CDS) of human *MPV17* was picked by PCR from HEK293T total cDNA whereas the presequence of the Ornithine Transcarbamylase was picked by PCR from the previously generated preOTC-DHFR plasmid (Meurant et al., 2023). Similarly, the CDS of the human *TWNK* gene was picked by PCR using the Twinkle-APEX2-V5 plasmid, a generous gift from Dr. Alice Ting (Han et al., 2017) (Addgene plasmid # 129705; http://n2t.net/addgene:129705; RRID:Addgene_129705). All sequences were cloned in a pDONR201 donor plasmid, kindly provided by Dr. Xavier DeBolle, using the Gateway™ BP Clonase™ II Enzyme mix (ThermoFisher Scientific). The sequences were subsequently cloned in the destination vector MAC-tag-C, a gift from Markku Varjosalo (Liu et al., 2018) (Addgene plasmid # 108077; http://n2t.net/addgene:108077; RRID:Addgene_108077), using the Gateway™ LR Clonase™ II Enzyme mix (ThermoFisher Scientific). The final vectors sequences were validated by Sanger sequencing (Eurofins Discovery).

The MPV17-HA plasmid was obtained by cloning the *MPV17* CDS into the pKH3, a gift from Ian Macara (Mattingly & Macara, 1996) (Addgene plasmid # 12555; http://n2t.net/addgene:12555; RRID:Addgene_12555). Briefly, the *MPV17* CDS was picked by PCR using a home-made pMPV17-pcDNA3.1 plasmid as a template. The insert was then cloned in the pKH3 plasmid using classical restriction-ligation with HindIII and XbaI restriction enzymes (Promega, Madison, USA) and the final vector sequence was validated by Sanger sequencing (Eurofins Discovery).

The mito-roGFP2-ORP1 reporter was cloned from the pMOS015E: Entry vector for: mito-roGFP2-Orp1 H2O2 oxidation sensor (mitochondrial), a gift from Adam Cohen (Werley et al., 2020) (Addgene plasmid # 163076; http://n2t.net/addgene:163076; RRID:Addgene_163076). The sequence of the construct was cloned into pKH3 plasmid for its use in transfection experiment for mammalian expression. Therefore, the STOP codon of the construct was kept. The cloning was done by classical restriction-ligation using SalI and XbaI restriction enzymes (Promega). The final vector sequence was validated by Sanger sequencing (Eurofins Discovery).

### 2.4 Site-Directed Mutagenesis

Site directed mutagenesis was performed using the Q5® Site-Directed Mutagenesis Kit according to the manufacturer’s instruction (New England BioLabs Inc. (NEB), Ipswich, USA). The mutagenesis primers for the addition of the STOP codon at the end of the *MPV17* gene from MPV17-HA plasmid (F: 5’-TAATCTAGAATGTACCCATACGATGTTC-3’; R: 5’-GAGCCGATGTGCCTTCCA-3’) were generated using the NEBaseChanger bioinformatic tool (NEB). The mutation was confirmed and the plasmids were validated using Sanger sequencing (Eurofins Discovery).

### 2.5 Transfection of MPV17 constructs

HEK293T cells were seeded at 30 000/45 000 cells/cm^2^ and transfected 24 h after with plasmids (pcDNA3.1-GFP, MPV17-HA or MPV17) preincubated with XTremeGENE HP Transfection Reagent (Roche, Basel, Switzerland) for 30 min at room temperature in Opti-MEM I serum-free medium (ThermoFisher Scientific) (ratio: 1 μg of DNA per 3 μL of transfecting reagent). After 24 h, the medium was replaced with fresh medium. After 72 h of plasmid expression, cells were harvested for DNA extraction.

### 2.6 DNA extraction

Cells were seeded at 30 000 cells/cm^2^, harvested 24 h later or differentially treated for several days depending on the experiment and then were harvested for DNA extraction. The DNA was extracted using the Wizard Genomic DNA Purification Kit (Promega, Madison, USA), according to the manufacturer’s instruction. The DNA was then diluted to 0.67 ng/µL and mtDNA content was assessed by qPCR using primers targeting the mitochondria-encoded genes NADH dehydrogenase subunit 1 (*ND-1*) and NADH dehydrogenase subunit 2 (*ND-2*) and normalized to the nuclear genomic DNA by using primers targeting TATA-Binding Protein (*TBP*) and *BECLIN* genes. qPCR was performed using the Gotaq qPCR Master Mix (Promega) with the StepOne Real Time PCR (Applied Biosystems, Waltham, USA) or ViiA 7 system (Applied Biosystems). The 2^-ΔΔCt^ method was used to express the normalized abundance of the mitochondrial genes in the WT cells relatively to their normalized abundance in KO cells.

### 2.7 mtDNA recovery assay

Cells were seeded at 30 000 cells/cm^2^ and treated 24 h later with 10 nM ethidium bromide, supplemented with 50 μg/mL uridine and 1 mM pyruvate for three days with a daily renewal. Cells were then treated with or without 100 µM dNTPs mix (Promega) and were harvested 0, 2, 3, 4 and 5 days after the mtDNA-induced depletion, as described in Figure 1B. DNA extraction was then performed and the mtDNA levels were measured by qPCR. In order to quantify the difference in the recovery profile between the conditions, an exponential regression was calculated and the slope of the regressions were compared.

**Figure 1.**
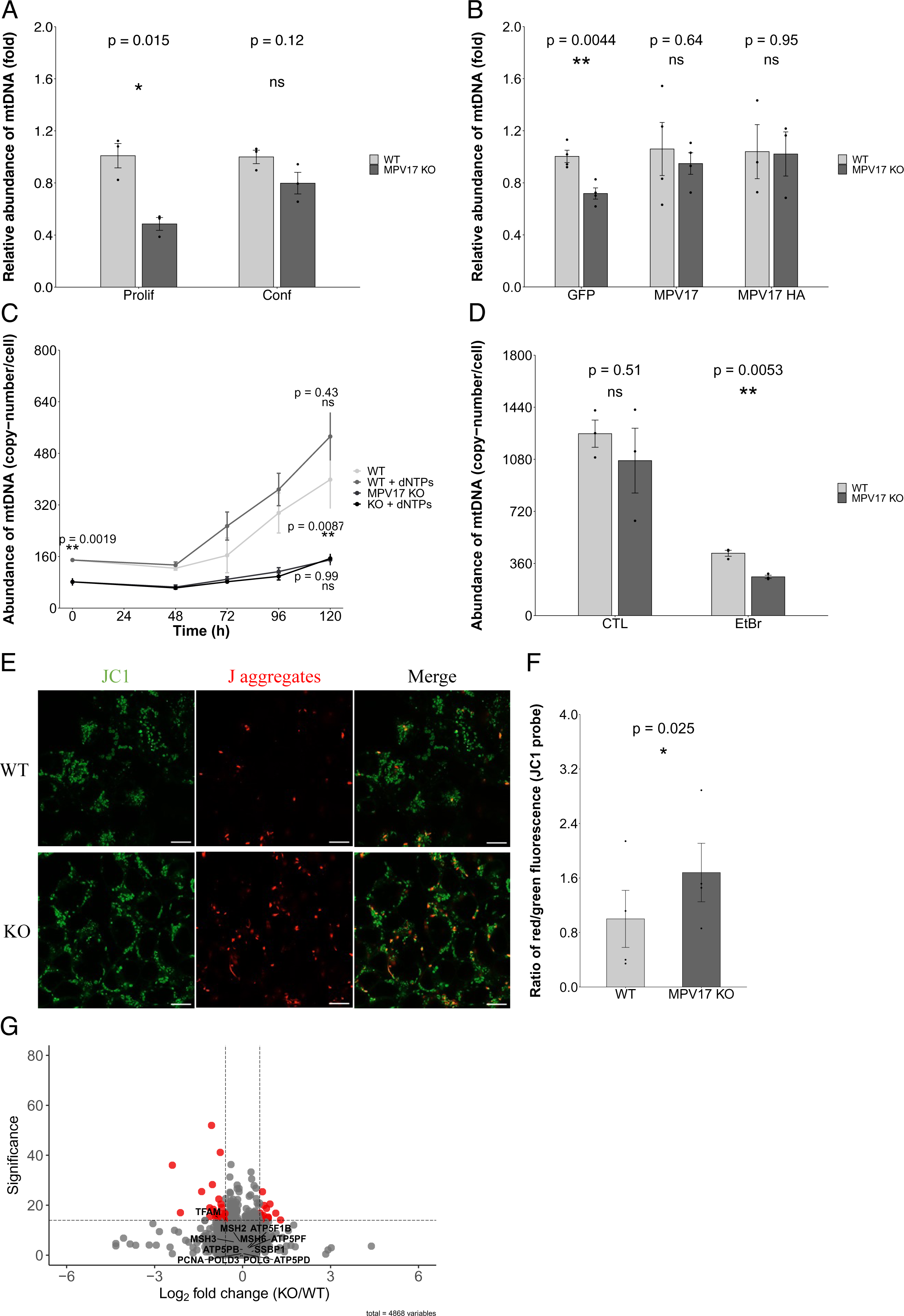
MPV17 KO cell lines characterization. (**A**) HEK293T cell lines WT and KO for MPV17 were generated using CRISPR-Cas9 technology and selected after limiting dilution and clonal expansion. Cells were seeded at low confluency (30 000 cells/cm^2^, proliferating cells) or high confluency (120 000 cells/cm^2^, confluent cells) and harvested 24 h after for DNA extraction. mtDNA abundance was measured using qPCR with primers targeting the mitochondrial genes ND1 and ND2 and was normalized using the abundance of the nuclear genes Beclin and TBP. Data are expressed relative to WT cells condition-wise and an unpaired t-test was performed for each condition (n = 3 using at least two different WT and KO clones). Data are represented as mean ± SEM. * p < 0.05. (**B**) HEK293T^MPV17+/+^ and HEK293T^MPV17-/-^ cells were seeded, transfected 24 h after with pcDNA3.1-GFP, MPV17-HA or untagged MPV17 plasmid and were harvested 72 h after for DNA extraction. To be able to observe the depletion phenotype at confluency, 10 nM of ethidium bromide supplemented with 50 µg/mL uridine and 1 mM pyruvate were added for the last 24 h prior DNA extraction. mtDNA abundance was measured using qPCR. Data are expressed relative to WT cells for each plasmid and an unpaired t-test was performed (n = 4 for GFP and MPV17 conditions and n = 3 for MPV17-HA condition using two different clones for WT and one clone for KO cells). Data are represented as mean ± SEM. ** p < 0.01. (**C**) HEK293T^MPV17+/+^ and HEK293T^MPV17-/-^ were seeded and treated 24 h after with 10 nM ethidium bromide supplemented with 50 µg/mL of uridine and 1 mM sodium pyruvate for 72 h. The medium was then refreshed with medium containing either 0 or 100 µM of dNTPs mix and cells were harvested at 0, 48, 72, 96, 120 h after for DNA extraction. mtDNA abundance was measured using qPCR. An ANOVA followed by a Tukey HSD post-hoc test was performed on the 0 and 120 h timepoints (n = 3 independent experiments). (**D**) HEK293T^MPV17+/+^ and HEK293T^MPV17-/-^ cells were seeded and treated 72 h after with 0 (CTL) or 10 nM ethidium bromide (EtBr) supplemented with 50 µg/mL of uridine and 1 mM sodium pyruvate for 24 h prior DNA extraction. mtDNA abundance was measured using qPCR. An unpaired t-test was performed (n = 3 using at least two different clones). Data are represented as mean ± SEM. ** p < 0.01. (**E-F**) HEK293T^MPV17+/+^ and HEK293T^MPV17-/-^ cells were seeded and stained 24 h later with 10 µg/mL of JC1 probe for 30 min at 37°C. Cells were then imaged using a Zeiss LSM900 confocal microscope. Scale bar = 10 µm. Quantification of the red/green ratio of fluorescence was performed using Fiji (ImageJ2) software (**F**). A paired t-test was performed and quantifications were done on 75-100 cells per replicate (n = 4 independent experiments). Data are represented as mean ± SEM. * p < 0.05. (**G**) HEK293T^MPV17+/+^ and HEK293T^MPV17-/-^ cells were seeded and harvested 24 h after for protein extraction. An amount of 10 µg of proteins were digested by FASP and the digested peptides were then analyzed using a nanoLC-ESI-MS/MS tims TOF Pro coupled with an UHPLC nanoElute. The identification and label-free quantifications were done using PEAKS Studio 11 software. A 1% FDR and a single peptide detection threshold were set. Data are represented as a measure of enrichment of proteins in MPV17 KO cells compared to WT ones (n = 3). Different proteins of interest are displayed on the volcano plot and a 15-significance threshold and 1.5-fold-change threshold were set.

### 2.8 Luciferase reporter assay

Cells were seeded at 30 000 cells/cm^2^ and were transfected 24 h later with NRF1, CREB, YY1 or TFAM luciferase reporters in combination with pCMV-βGal reporter, each preincubated with XTremeGENE HP Transfection Reagent for 30 min at room temperature in Opti-MEM I serum-free medium (ratio: 1 μg of DNA per 3 μL of transfecting reagent). 48 h after, cells were lysed in Reporter Lysis Buffer (Promega) and processed according to manufacturer’s instruction. Briefly, the lysates were frozen at -80°C for 15 min, shaken for 5 min at room temperature and transferred to new tubes. The lysates were then vortexed for 15 seconds and centrifuged at 16.000× g for 30 seconds. Then each supernatant was transferred into a new tube before the revelation with either Luciferase Assay Reagent (Promega) or with β galactosidase (βGal) buffer (2 mM MgCl_2_.6H_2_O, 15 µM of O-nitrophenyl-beta-D-galacto pyranoside (ONPG), 0.7 % of β-mercaptoethanol and water), following manufacturer’s instructions. To assess the luciferase reporter activity, the relative light units (RLU) value was divided by the 405 nm optical density value obtained from βGal measurement.

### 2.9 Western Blotting (WB)

Western blotting was performed as previously described (Meurant et al., 2023). Briefly, HEK293T cells were scraped and lysed in homemade radioimmunoprecipitation assay (RIPA) lysis buffer (10 mM Tris-HCl; pH 8.0, 1 mM EDTA, 0.5 mM EGTA, 1% Triton X-100, 0.1% Sodium Deoxycholate, 0.1% SDS, 140 mM NaCl) complemented with SuperNuclease (25 U/µL, Bio-connect), complete Protease Inhibitor Cocktail (Roche) and homemade Phosphatase Inhibitor Cocktail (25 mM Na_3_VO_4_, 250 mM 4-nitrophenylphosphate, 250 mM di-Sodium β-glycerophosphate pentahydrate and 125 mM NaF) and maintained on ice. Mixing with a thermomixer was performed for 10 min at 4 °C and 1400 RPM. Cell lysates were then centrifuged for 10 min at 16.000× g at 4 °C to collect supernatants. Sample protein content was determined with the Pierce detection assay (Pierce 660 nm Protein Assay Reagent (ThermoFisher Scientific), complemented with Ionic Detergent Compatibility Reagent (IDCR) (ThermoFisher Scientific) according to the manufacturer’s instructions.

An equivalent of 15-20 μg of proteins was prepared in Western blot loading buffer (30 mM Tris; pH 6.8, 1.2% SDS, 3% β-mercaptoethanol, 5% glycerol and 150 µM bromophenol blue), boiled for 5 min at 98 °C and resolved on home-made polyacrylamide gel (SDS-PAGE). The Color Protein Standard Broad Range (10–250 kDa) (New England BioLabs Inc.) was used as protein molecular weight ladder. Protein resolution was performed at 150–200 V (400 mA and 100 W). Proteins were then transferred on a PolyVinyliDene Fluoride (PVDF) membrane (Immobilon-P, Merck Millipore, Burlington, USA) using liquid transfer with 20 % methanol at 100V, at 4 °C for 2 h. The PVDF membrane was then incubated for 1 h with the Intercept Blocking Buffer (TBS) (Li-cor Biosciences, Lincoln, USA) blocking solution at room temperature. The primary antibody solutions were prepared in the blocking solution containing 0.1 % Tween-20 (Roth) and incubated overnight with the membrane on a rocker at 4 °C. The primary antibodies used were rabbit polyclonal Biotin (ab53494, Abcam) (Wang et al., 2021); mouse monoclonal BirA (NBP2-59939, Novus Biologicals, Abingdon, UK) (Creed et al., 2020); rabbit polyclonal CHCHD3/MIC19 (25625-1-AP, Proteintech, Rosemont, USA) (Kondadi et al., 2020); rabbit polyclonal Mitofilin/MIC60 (10179-1-AP, Proteintech) (Sohn et al., 2023); rabbit polyclonal PPIF (18466-1-AP, Proteintech) (Jia et al., 2023); and mouse monoclonal HA-tag (2367, Cell Signaling Technology (CST), Danvers, USA) (Bergant et al., 2023). The secondary antibody solution was prepared in the blocking solution with 0.1% Tween-20 and the following: anti-rabbit goat polyclonal antibody (IR Dye 800CW) (926-32211, Li-cor Biosciences); anti-rabbit goat polyclonal antibody (IR Dye 680RD) (926-68071, Li-cor Biosciences) and anti-mouse goat polyclonal antibody (IR Dye 680RD) (926-68070, Li-cor Biosciences). Membrane fluorescence was detected using the Odyssey Li-cor Scanner (Li-cor Biosciences).

### 2.10 Immunoprecipitation

Immunoprecipitation was performed as previously described (Meurant et al., 2023). HEK293T cells were scraped in RIPA buffer and the lysate was incubated for 30 min on a rotating wheel at 4 °C before a centrifugation of 10 min at 16.000× g at 4 °C. Sample protein content was determined with the Pierce detection assay and 200-800 µg of proteins were loaded on Dynabeads Protein G for Immunoprecipitation (ThermoFisher Scientific); the beads were previously incubated for 10 min on a wheel at room temperature with PBS-Tween-20 0.01% supplemented or not with 2.5-5 µg of MIC60, MIC19, ACBD5 or PPIF antibody. 5 % of total protein amount used for immunoprecipitation was saved for input. Protein lysates were incubated with the beads on a wheel at 4 °C for 16 h. The beads were then rinsed three times with NETN buffer (0.5 % NP40, 1 mM EDTA, 50 mM Tris-HCl; pH 8, 100 mM NaCl), twice with ETN (1 mM EDTA, 50 mM Tris-HCl; pH 8, 100 mM NaCl) and once with ddH_2_O. Beads were then resuspended either in Western blot loading buffer and boiled for 5 min at 98 °C before resolution by SDS-PAGE or were resuspended in 20 mM Tris-HCl (pH 8) before digestion and mass spectrometry analyses.

### 2.11 Proximity labelling assay (BioID)

HEK293T cells were seeded at 30 000 cells/cm^2^ and transfected 24 h after with 5 µg/T75 flask of either preOTC-BirA*, MPV17-BirA* or TWNK-BirA* plasmids. The medium was refreshed 24 h later and 48 h after transfection, cells were incubated for 6 or 18 h with medium supplemented with 50 µM biotin. The medium was then changed to remove biotin for 3 h before cell lysis. Cells were harvested, resuspended in RIPA buffer and incubated for 45 min on a thermo-mixer (Eppendorf) at 4 °C and 1000 RPM (Round Per Minute). The lysates were then sonicated three times for 10 s at 70 % amplitude, frequency 1, and centrifuged for 15 min at 16.000× g at 4 °C to collect supernatants. Sample protein content was determined with the Pierce detection assay and a maximum amount of proteins (2.3-4 mg) was loaded on 1 mg of Dynabeads M-280 Streptavidin (ThermoFisher Scientific), previously equilibrated with RIPA buffer. An equivalent of 0.5 % of total proteins was saved as the input for Western blot analysis. Protein lysates were incubated with the beads on a rotating wheel at 4 °C for 6-18 h. The beads were then washed according to Le Sage and colleagues (Le Sage et al., 2016). Briefly, proteins were successively washed with washing buffer 1 (2 % SDS in sterile deionized water), washing buffer 2 (50 mM HEPES; pH 7.5, 500 mM NaCl, 1 mM EDTA, 0.1 % sodium deoxycholate, 1 % Triton X-100) and washing buffer 3 (10 mM Tris-HCl pH 7.4, 250 mM LiCl, 1 mM EDTA, 0.1 % sodium deoxycholate, 1 % NP-40). Finally, the beads were washed twice with 20 mM Tris-HCl (pH 8) and resuspended in 20 mM Tris-HCl (pH 8). An amount corresponding to 5 % of the pull-down volume was saved for Western blot analyses and 1 µg of trypsin gold (Promega) was then added to the beads for 16 h at room temperature in a thermomixer at 600 RPM. Supernatant was then transferred to a new vial and 500 ng of trypsin was added to the beads followed by a 3 h incubation at room temperature in a thermomixer at 400 RPM. Trypsin was then inactivated with 2 % formic acid (Biosolve). The digested peptides were then desalted using Pierce C18 Spin Tips & Columns system according to the manufacturer’s instructions (ThermoFisher Scientific). Three independent biological replicates for each condition were analyzed by mass spectrometry.

### 2.12 Mass spectrometry analyses

For the mass spectrometry analyses performed on total protein lysates, 10 µg of proteins were digested by Filter Aided Sample Preparation (FASP) as previously described (Nicolas et al., 2023). All mass spectrometry analyses were performed as previously described (Meurant et al., 2023). Briefly, digested proteins were analyzed using nanoLC-ESI-MS/MS tims TOF Pro (Bruker, Billerica, USA) coupled with an UHPLC nanoElute (Bruker). Peptides were analyzed by nanoUHPLC (nanoElute, Bruker) on a 75 μm ID, 25 cm C18 column with integrated CaptiveSpray insert (AuroraIonOpticks, Victoria, Australia) at a flow rate of 200 nL/min, at 50 °C. LC mobile phase A was 0.1% formic acid (v/v) in water and B was formic acid 0.1% (v/v) in acetonitrile. Peptides were injected, and the gradient was increased from 2% B to 15% in 40 min, from 15% B to 25% in 15 min, from 25% B to 37% in 10 min and from 37% B to 95% in 5 min. Data acquisition on the timsTOF Pro was carried out using Hystar 5.1 and timsControl 2.0. Data were acquired using 160 ms TIMS accumulation time, mobility (1/K0) range from 0.75 to 1.42 Vs/cm^2^. Mass spectrometry analysis was performed using the parallel accumulation serial fragmentation (PASEF) acquisition method (Meier et al., 2018). Cycles of one MS spectra followed by six PASEF MSMS spectra in a total duration of 1.16 s were carried out. Two injections per sample were performed.

Data analysis was carried out using PEAKS Studio 11 X Pro with ion mobility module and Q module for label-free quantification (Bioinformatics Solutions Inc., Waterloo, Canada). PEAKS software was used to identify the proteins with 15 ppm as parent mass error tolerance and 0.05 Da as fragment mass error tolerance. The oxidation of methionine, biotinylation of lysine and acetylation (N-term) were allowed as variable modifications. Trypsin was used as a digestion enzyme and the maximum number of missed cleavages per peptide was set at two. The protein library used was Homo Sapiens with isoforms from UNIREF 100 (195195 sequences). Peptide spectrum matches and protein identification was normalized to less than 1.0 % false discovery rate.

The label-free quantitation (LFQ) method is based on the expectation–maximization algorithm on the extracted ion chromatograms of the three most abundant unique peptides of a protein to calculate the area under the curve (Lin et al., 2013). Mass error and ion mobility tolerance were set, respectively, to 15 ppm and 0.08 1/k0 for the quantitation. Peptide quality score was set to ≥ 3 and the protein significance score threshold was set to 15. The significance score is calculated as the −10log10 of the significance testing p-value (0.05), with the ANOVA used as the significance testing method. Total ion current was used for the normalization of each extracted ion current.

The exported PEAKS data of label-free quantification were sorted and represented using R v4.3.0 software with the “EnhancedVolcano” package (Blighe et al., 2023). The fold change cutoff was set at 2 or 1.5 and the p-value cutoff was set at 10 × 10^−15^ (significance of 15).

For MPV17 KO cell lines screening and for MIC60 immunoprecipitation and mass spectrometry analysis, data analysis was performed using Mascot 2.8.1 (Matrix Science, London, UK; version 2.8.1). Briefly, Tandem mass spectra were extracted, charge state deconvoluted and deisotoped by Data analysis (Bruker) version 5.3. All MS/MS samples were analyzed using Mascot. Mascot was set up to search the Homo Sapiens with isoforms from UNIREF 100 (195195 sequences) assuming the digestion enzyme trypsin. Mascot was searched with a fragment ion mass tolerance of 0.050 Da and a parent ion tolerance of 15 PPM. Carbamidomethyl of cysteine was specified in Mascot as fixed modifications. Oxidation of methionine and acetyl of the n-terminus were specified in Mascot as variable modifications. For protein identification, Scaffold (version Scaffold_5.0.0, Proteome Software Inc., Portland, OR) was used to validate MS/MS based peptide and protein identifications. Peptide identifications were accepted if they could be established at greater than 96.0% probability to achieve an FDR less than 1.0% by the Scaffold Local FDR algorithm. Protein identifications were accepted if they could be established at greater than 5.0% probability to achieve an FDR less than 1.0% and contained at least 2 identified peptides. Protein probabilities were assigned by the Protein Prophet algorithm (Nesvizhskii, Al et al Anal. Chem. 2003;75(17):4646-58). Proteins that contained similar peptides and could not be differentiated based on MS/MS analysis alone were grouped to satisfy the principles of parsimony. Proteins sharing significant peptide evidence were grouped into clusters.

The mass spectrometry proteomics data were deposited to the ProteomeXchange Consortium via the PRIDE (Perez-Riverol et al., 2022) partner repository, with the dataset identifier PXD051217 and 10.6019/PXD051217. Data can be accessed using the username: and the password: SazcOxgX.

### 2.13 Mitophagy and autophagy assessment

Cells were seeded at 45 000 cells/cm^2^ on coverslips coated with poly-L-lysine (Sigma-Aldrich). After 24 h, cells were transfected with pFis1-mCherry-GFP (Allen et al., 2013) or pLC3B-mCherry-GFP (Pankiv et al., 2007), to monitor mitophagy and autophagy, respectively. Cells were first preincubated with XTremeGENE HP Transfection Reagent (Roche) for 30 min at room temperature in Opti-MEM I serum-free medium (ratio: 1 μg of DNA per 3 μL of transfecting reagent). After 24 h, cells were washed twice with PBS, fixed with 4 % of pre-warmed paraformaldehyde during a 30-min incubation at room temperature and rinsed again twice with PBS. Fixed cells were then permeabilized for 15 min, with PBS-Triton (0.6 %). A 30-min incubation of PBS supplemented with 1 µg/mL of 4′,6-diamidino2-phenylindole (DAPI) (Sigma-Aldrich) was performed. Mounting of the slides was performed with FluoromountG (ThermoFisher Scientific). Samples were imaged using a Zeiss LSM900 confocal microscope, with an Airyscan detector and a 63x objective (final resolution of 120 nm) (Zeiss, Oberkochen, Germany). To assess mitophagy and autophagy, the properties of the probe were used: in acidic environment the Green Fluorescent protein is quenched, and the only emission spectrum observed is mCherry (red), while in physiological pH, both mCherry and GFP emission spectra are detected (yellow). Mitophagy and autophagy were thus calculated as the mean number of red puncta per cells and each red puncta was counted manually using Fiji.

### 2.14 Immunofluorescence analyses

Cells were seeded at 30 000/45 000 cells/cm^2^ on coverslips coated with poly-L-lysine and exposed to different experimental conditions. They were then washed three times with PBS, fixed with 4 % paraformaldehyde and rinsed again three times with PBS. Fixed cells were then permeabilized for 5 min with PBS-0.5 % Triton X-100. Cells were then incubated for 1 h in the blocking solution (PBS containing 2 % Bovine Serum Albumin (BSA), 5 % FBS and 0.2 % Triton X-100). Primary antibody solutions for immunostaining were prepared in blocking solution and incubated with fixed cells at 4 °C overnight. The primary antibodies used were mouse monoclonal BirA (NBP2-59939, Novus Biologicals); rabbit polyclonal TOM20 (ab186734, Abcam, Cambridge, UK); mouse monoclonal HA-tag (2367, CST); rabbit polyclonal DNA-PKcs (ab32566, Abcam); and mouse monoclonal mtHSP70 (ALX-804-077-R100, Enzo, New-York, USA).

The next day, the cells were rinsed twice for 10 min with blocking solution. Cells were then incubated with secondary antibody in blocking solution supplemented with 1 µg/mL of DAPI intercalating agent for 1 h at room temperature in the dark. Secondary antibodies used were goat polyclonal anti-rabbit (Alexa Fluor 488 nm) (A-11008, ThermoFisher Scientific); goat polyclonal anti-mouse (Alexa Fluor 568 nm) (A-11031, ThermoFisher Scientific); and for biotin staining, the streptavidin-Alexa488 probe was used (S32354, ThermoFisher Scientific). Cells were then rinsed twice with blocking solution for 10 min and then left in PBS. The mounting of the slides was performed with warmed Mowiol (Sigma-Aldrich) or Fluoromount G. Images were acquired on either Leica SP5 Confocal Microscope with a 63x objective (final resolution of 200 nm), using the LAS-AF Lite v4.0 software (Leica microsystem, Wetzlar, Germany), or a Zeiss LSM900 confocal microscope with the GaAsP-PMT or Airyscan detector and with a 63x objective, using the ZEN v3.7 software (Zeiss).

### 2.15 Stimulated-Emission-Depletion (STED) microscopy

Cells were seeded at 60 000 cells/cm^2^ and incubated 24 h after with 100 nM of Live Orange probe (Abberior, Goettingen, Germany) for 30 min at 37 °C and 5 % CO_2_. The medium was then refreshed and images were acquired 10 min after on an INFINITY (Abberior) or STEDYCON (Abberior) microscope with a 63x objective (final resolution of ∼50 nm), using either INFINITY or STEDYCON browser-based softwares.

### 2.16 Mitochondrial potential measurement

Cells were seeded at 30 000 cells/cm^2^ and incubated 24 h after with 10 µg/mL of JC1 probe (ThermoFisher Scientific) for 30 min at 37 °C and 5 % CO_2_. The medium was then refreshed and confocal images were acquired 10 min after on a Zeiss LSM900 confocal microscope with a 63x objective. To determine the mitochondrial potential, the ratio of the J-aggregates (red fluorescence) signal over the monomeric JC1 probe (green fluorescence) signal was calculated using Fiji software.

### 2.17 Mitochondrial calcium measurement

Cells were seeded at 30 000 cells/cm^2^ and were treated 6 h later with 6 µM of the calcium-sensitive Rhod2AM probe (ThermoFisher Scientific) for 1 h before refreshing the cell culture medium. The probe was incubated for 18 h for mitochondrial accumulation as described previously (Zhu et al., 2000) and the cells were then incubated for 10 min with 10 µg/mL of Rhodamine 123 (ThermoFisher Scientific) for mitochondrial staining. The medium was then refreshed and confocal images were acquired on a Zeiss LSM900 confocal microscope with a 63x objective. The mitochondrial calcium content was then quantified using Fiji software.

### 2.18 mPTP opening assessment using mitochondrial calcium tracking

Cells were seeded at 30 000 cells/cm^2^ and transfected 24 h after with the calcium reporter pCAG mito-GCaMP5G, a generous gift from Franck Polleux (Kwon et al., 2016) (Addgene plasmid # 105009; http://n2t.net/addgene:105009; RRID:Addgene_105009), preincubated with XTremeGENE HP Transfection Reagent (Roche) for 30 min at room temperature in Opti-MEM I serum-free medium (ratio: 1 μg of DNA per 2 μL of transfecting reagent). 48 h after, cells were treated with 500 nM of ionomycin (Sigma-Aldrich) in combination or not with 5 µM cyclosporin A (Selleck Chemicals). The variation of the reporter fluorescence intensity before and after ionomycin addition was measured every min for 30 min on a Zeiss LSM900 confocal microscope with 63x objective. The fluorescence intensity of each cell was quantified overtime using Fiji software and was expressed relatively to the fluorescence of the construct before the stress (T = 0 min). The maximal value of fluorescence (expressed in fold compared to T = 0 min) reached in each cell before the loss of fluorescence due to mPTP opening was used as a measure of the maximal calcium threshold triggering mPTP opening for each cell and was compared between the different conditions.

### 2.19 Mitochondrial ROS measurement

Cells were seeded and transfected 24 h after with the mitochondrial H_2_O_2_ reporter pmito-roGFP2-ORP1, preincubated with XTremeGENE HP Transfection Reagent (Roche) for 30 min at room temperature in Opti-MEM I serum-free medium (ratio: 1 μg of DNA per 2 μL of transfecting reagent). 48 h after, confocal images were acquired, with the sequential excitation of the ratiometric construct at first 405 nm and then 488 nm, on a Zeiss LSM900 confocal microscope with a 63x objective. The mitochondrial ROS content was then quantified, using Fiji software, by dividing the fluorescence measured following 405 nm excitation by the fluorescence acquired following 488 nm excitation.

### 2.20 Transmission Electron Microscopy (TEM)

The samples used for transmission electron microscopy were processed using standard protocols. Cells were fixed in 2.5 % electron microscopy-grade glutaraldehyde (LFG Distribution, Sainte Consorce, France) in 0.1□M phosphate buffer pH 7.4 for 16□h at 4□°C. Samples were rinsed with 0.1□M phosphate buffer and post-fixed by incubation with 1 % osmium tetroxide and 1 % potassium ferricyanide in ddH_2_O for 60□min at room temperature. Samples were then rinsed with distilled water, dehydrated in graded series of ethanol solutions and finally embedded in Epon at 60□°C for 48□h. Embedded samples were cut into 60□nm-thick sections, which were contrast-stained with 3 % uranyl acetate in ethanol 50° for 15□min and then observed under a JEOL JEM 1400 transmission electron microscope (JEOL, Tokyo, Japan) operating at 120□keV and equipped with a Gatan Orius digital camera (Gatan, Pleasanton, USA). Analysis and measurements were performed on 5-10 cells per replicate. Mitochondria were identified by the presence of a double membrane with the inner membrane shaping folding in the matrix space.

### 2.21 Image analysis

All fluorescence image analyses were done using Fiji (ImageJ2) software and at least on three independent biological replicates with two-three different WT and KO cell lines and on 75-100 cells per replicate except for the autophagy and mitophagy measurement where 20-50 transfected cells were used and the mPTP opening experiment where only 3-4 cells were analyzed per replicate. For the TEM analyses, MDV events were defined as budding of the mitochondrial outer membrane and were manually counted and normalized over the number of mitochondria, their average area was also calculated.

For the quantification of the mitochondrial calcium content, a home-made macro was prepared. Briefly, the Rhodamine 123 signal was used as a mitochondrial mask and the mean gray value of the Rhod2AM fluorescence was measured inside this mask and was then expressed relatively to the WT control condition. For the fluorescence intensity quantification of the mito-CaMP5G, a home-made macro was prepared. For each stack, containing the untreated + the 31 micrographs acquired after ionomycin treatment, responding cells were manually delimited and the total fluorescence value was measured and normalized by the area of the signal surface for each slice. Each normalized fluorescence value was then expressed relatively to the fluorescence of the corresponding untreated (T = 0 min) cell.

### 2.22 Data analyses

All data and statistical analyses were performed using the statistical programming language R (http://www.rproject.org/ accessed on 6 October 2022). Paired or unpaired t-tests were done for 2-by-2 comparisons. For the recovery experiment and mPTP opening experiment, an ANOVA3 and ANOVA followed by TukeyHSD post hoc test for multiple comparisons were performed after data normality verification. An Over Representation Analysis was performed on the MPV17 BioID datasets using the DAVID platform (Dennis et al., 2003) and using as input data all the proteins showing ≥ 2-fold-change enrichment compared to the control condition. In addition, an interaction network analysis was performed also on those data using the STRING platform (von Mering et al., 2003). A Gene Set Enrichment Analysis was performed on the BioID whole datasets using R and the “ClusterProfiler” package (Wu et al., 2021). The representations of the GSEA data were done using the “enrichplot” (Yu, 2021) and “pathview” (Luo & Brouwer, 2013) packages. The analyzed and sorted BioID and proteomic data are available as Dataset EV (1-4).

### 2.23 Computational analysis

Protein modeling (Jumper et al., 2021) and protein multimer prediction (Evans et al., 2022) have been made using the AlphaFold Collab notebook (v1.5.2), available on GitHub Repository. The heatmap of the predicted interactions were visualized using the Predicted Aligned Error (PAE) viewer (Elfmann & Stülke, 2023) after loading the Alphafold multimer prediction. All structural analyses have been made with ChimeraX (v1.6) Software (Meng et al., 2023). Alignment sequencing has been performed with the ConSurf server (Ashkenazy et al., 2016). According to them, 8213 homologs were collected from the UniRef90 database (Suzek et al., 2015) using HMMER v3.4 software (http://hmmer.org/). Of these, 1503 homologs passed the thresholds (min/max similarity, coverage, etc.), of which 1249 were unique to the CD-HIT. The calculations were performed on 150 results (including the query), sampled from the unique results. For the MPV17 channel modeling, the ChExVis tool was used (Masood et al., 2015). The PyMOL (v2.5) software was used for the measurement of conformational changes of MPV17 structure upon mutational load through Root Mean Square Deviation (RMSD) calculation.

## 3. DATA AVAILABILITY

The datasets described below which were produced in this study are available in the following database: PRIDE PDX (https://www.proteomexchange.org/; Accession: PXD051217 and DOI: 10.6019/PXD051217)

- Proteomic data for MPV17 detection (WT-KO)
- Protein interaction MIC60 IP-MS data (WT-KO)
- Proteomic data (KO/WT comparison)
- Protein interaction PLA-MS data (MPV17-BirA*/preOTC-BirA)
- Protein interaction PLA-MS data (MPV17-BirA*/Untransfected)
- Protein interaction PLA-MS data (TWNK-BirA* in MPV17 KO/WT)

## 4. RESULTS

### 4.1 Proliferating MPV17 KO cells display mtDNA depletion insensitive to dNTPs supplementation

To study the molecular function of MPV17 in human cells, we generated HEK293T MPV17 KO clones using CRISPR-Cas9 genome editing technology. The WT and the KO status of the clones was verified by Sanger sequencing analysis and MPV17 protein abundance assessment using mass spectrometry analysis (**Table EV1**), as the poor quality of MPV17 antibodies prevented Western blot analysis.

As the common MDDS feature is a mtDNA depletion in affected tissues, we assessed mtDNA level in the MPV17 KO cells and observed a significant depletion of mtDNA in proliferative cells, whereas the depletion was alleviated at confluency (**Fig.1A**). This phenotype was rescued after transfection by plasmids encoding either tagged or untagged version of MPV17, but not when using a plasmid encoding a cytosolic GFP (**Fig.1B**). The fact that this depletion was observed in proliferative cells is not in line with the hypothesis suggesting that MPV17 is involved in the purine mitochondrial salvage pathway (Dalla Rosa et al., 2016). Therefore, similarly to what was done previously (Dalla Rosa et al., 2016), we depleted mtDNA using ethidium bromide and we supplemented MPV17 WT and KO cells with dNTPs for five days. No effect of the dNTPs was observed on KO cells, while they led to slightly higher mtDNA level in WT cells, although not significant (**Fig.1C**). In addition, we observed that blocking mtDNA replication at confluency, using ethidium bromide, phenocopies the mtDNA depletion observed in proliferating cells (**Fig1.D**), suggesting that the mtDNA depletion in KO cells does not originate from a reduced replication but rather from another parameter, such as a putative higher degradation in KO cells.

We then assessed the mitochondrial membrane potential in MPV17 KO cells using the ratiometric JC1 probe (**Fig.1E**), and observed significantly more J aggregates witnessing a higher membrane potential in KO cells. This phenotype is thus not associated with a low energetic profile of the KO cells (**Fig.1F**). We further performed a comparative proteomic analysis of KO and WT cells (**Dataset EV1**) and disclosed only 9 and 21 proteins up and down-regulated respectively, when considering fold-change thresholds of 1.5 and 0.67 (**Fig.1G**).

### 4.2 Investigation of MPV17 proxisome

To gain insights in MPV17 proxisome, we used the BioID method (Kim et al., 2009) with the fusion of MPV17 to the modified biotin ligase BirA*. As controls, we used either untransfected cells or a modified BirA* targeted to the matrix, using the mitochondrial targeting sequence of the Ornithine Transcarbamylase (preOTC-BirA*) as depicted in **Figure 2A**. The mitochondrial localization of both BirA*-based constructs was verified by immunofluorescence and colocalization with TOM20 (**Fig EV.1A**). The biotinylation pattern of the constructs also displayed a mitochondrial localization, colocalizing with BirA* staining (**Fig.2B**). The biotinylating activity of the construct was also validated by Western blot analysis with specific biotinylating patterns for each construct, enriched following streptavidin-based pull-down (**Fig.2C**). While comparing the proteins enriched in the MPV17-BirA* condition compared to the preOTC-BirA* condition, 316 proteins were found with ≥ 2-fold-change (**Fig.2D**) (**Dataset EV2**). Similarly, in the MPV17-BirA*/Untransfected comparison, 359 proteins were found with ≥ 2-fold-change (**Fig.2E**) (**Dataset EV3**). In both comparisons, MPV17 was found as the most significantly enriched protein, as expected. An Over Representation Analysis (ORA) of cell component (CC) enrichment performed on all the proteins with a ≥ 2-fold-change value revealed a clear enrichment of mitochondria-related GO terms for both comparisons (**Fig EV.1B**). The high enrichment of the “mitochondrial inner membrane” GO term supports the localization of the MPV17-BirA* construct in the Inner Mitochondrial Membrane (IMM). Interestingly, in the MPV17-BirA*/Untransfected, the nuclear DNA Damage Response (DDR) protein MRE11 is only detected in the MPV17-BirA* condition (**Fig.2E**). Similarly, another DDR protein is enriched in the MPV17-BirA*/preOTC-BirA* comparison: the DNA-PKcs or PRKDC (**Fig.2D**) and its localization inside the mitochondria was verified by immunofluorescence in both WT and MPV17 KO cells (**Fig EV.1C-D**). However, no major difference was observed between both cell lines in term of PRKDC subcellular pattern or abundance (**Fig EV.1C)**.

**Figure 2.**
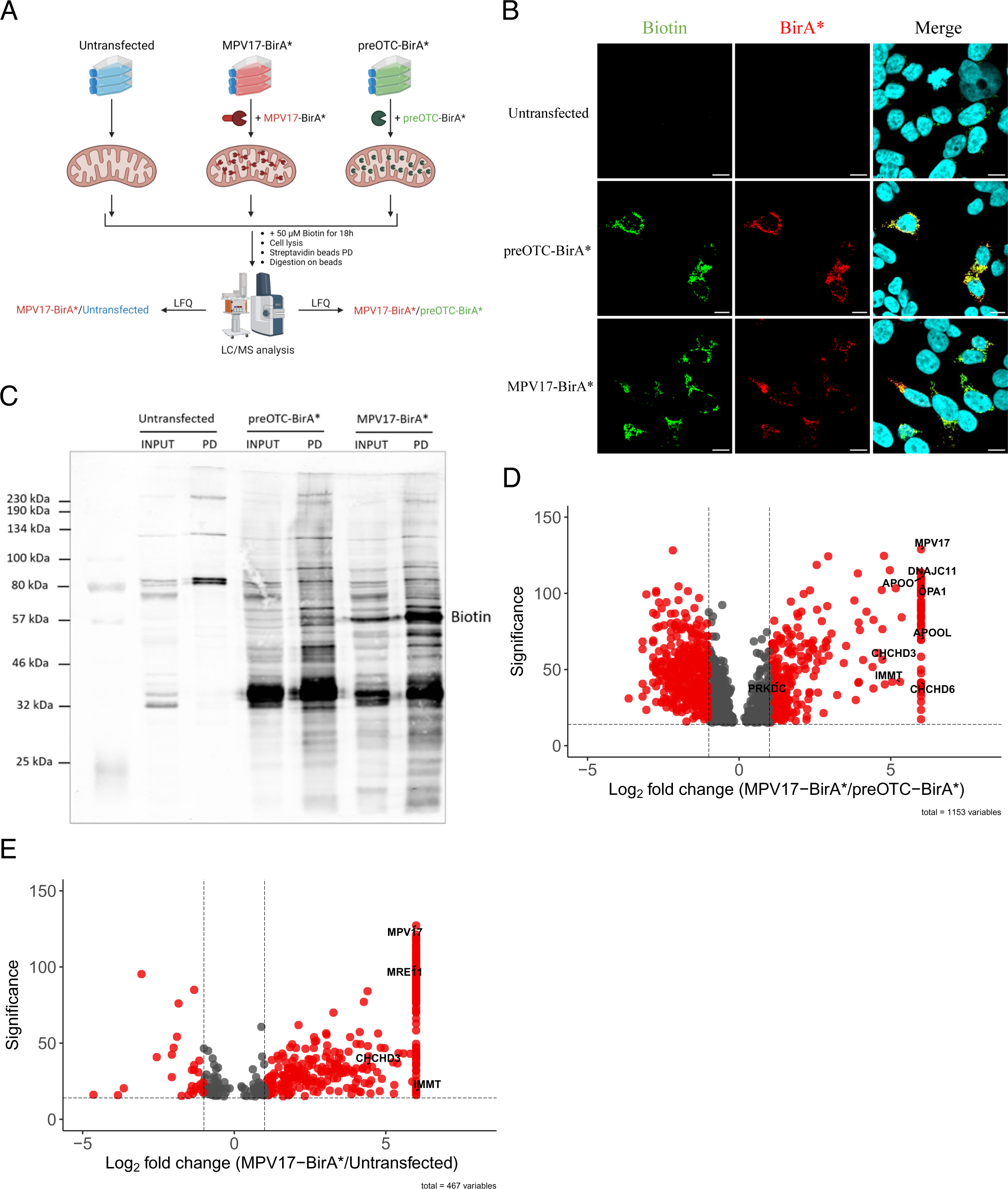
MPV17 proxisome identification using BioID. (**A**) Depicted strategy for MPV17-BioID. Briefly, HEK293T cells were seeded and transfected 24 h after with a plasmid coding either for MPV17-BirA* or a matrix-targeted biotin ligase (preOTC-BirA*) or were left untransfected. The cells were incubated 24 h after with 50 µM of biotin for 18 h before being harvested for protein extraction. The maximum amount of proteins (4 mg) was loaded on streptavidin beads and the biotinylated proteins were pulled-down for 16 h prior digestion on beads and mass spectrometry analysis, using a nanoLC-ESI-MS/MS tims TOF Pro coupled with an UHPLC nanoElute. The identification and label-free quantifications were done using PEAKS Studio 11 software. A 1% FDR and a single peptide detection threshold were set. The results were then represented as the enrichment in the MPV17-BirA* condition compared to either the preOTC-BirA* control or the untransfected one. The figure was generated on Biorender (https://app.biorender.com/). (**B**) HEK293T cells were seeded and transfected or not 24 h later with either MPV17-BirA* or preOTC-BirA* constructs. The cells were incubated 24 h after with 50 µM of biotin for 18 h before being fixed with 4 % PFA and labelled with anti-BirA* antibody and streptavidin-alexa488 probe to reveal biotinylated proteins. Nuclei were stained using DAPI. Micrographs were acquired on a Leica TCS SP5 confocal microscope. Scale bar = 10 µm. Representative experiment of three independent biological replicates. (**C**) HEK293T transfected with the constructs or not were processed according to **A**. After the pull-down, an equivalent of 0.5 % of total proteins (20 µg) was loaded on acrylamide gel as input along with 5 % of the pull-down (PD) for each experimental condition. Biotinylated proteins were revealed using specific antibody (n = 3). (**D-E**) Label-free quantifications showing the enrichment of the proteins in the MPV17-BirA* condition compared either to the preOTC-BirA* (**D**) or to the untransfected condition (**E**). Peptides detected in only one of the two experimental conditions were given an arbitrary 64-fold-change value. Proteins with a 15-significance threshold and 2-fold-change threshold are highlighted in red on the Volcano Plot. Different proteins of interest belonging to the MICOS complex (DNAGC11, OPA1, APOOL, IMMT, CHCHD3, CHCHD6), or related to DNA damage repair (PRKDC, MRE11), as well as MPV17, are displayed.

### 4.3 MPV17 interacts with the MICOS

Using the MPV17-BirA*/preOTC-BirA* comparison, the GO enrichment analysis revealed the “cristae formation” as the most significantly enriched biological process (BP) (**Fig.3A**) as well as the “MICOS complex”, among the most enriched Cellular Component (CC) (**Fig EV.2A**). Indeed, a high enrichment of core components of the MICOS complex (IMMT/MIC60, CHCHD3/MIC19, CHCHD6/MIC25, APOO/MIC26 and APOOL/MIC27) as well as components of the Membranes Inter-Bridging (MIB) complex (DNAJC11, OPA1), were observed in the MPV17-BirA* BioID (**Fig.2D**). Interestingly, a STRING network analysis also revealed the “mitochondrial crista junction” as the local network with the strongest significant interaction (**Fig.3B**) and is represented in the **Expanded View Figure 2B**, displaying the enriched proteins found in the proximity of MPV17 and part of this network of interaction. To confirm the interaction between MPV17 and the MICOS complex, we immunoprecipitated MIC60, the main subunit of the complex, and observed the co-immunoprecipitation of the MPV17-BirA* construct (**Fig.3C**). Similarly, the construct also co-immunoprecipitated with MIC19 and both MICOS proteins co-immunoprecipitated with each other, as expected (**Fig EV.2C**). The MICOS interaction was also validated endogenously using immunoprecipitation of MIC60 on mitochondrial fractions, followed by mass spectrometry analysis (**Table EV3**). In addition, *in silico* analyses using the multimer prediction tool of Alphafold (Evans et al., 2022; Jumper et al., 2021) highlighted a high confidence interaction of MPV17 with MIC60 displaying an interface Predicted Template Modeling (iPTM) score of 0.41 (**Fig.3D**). Of note, the 3D interaction modeling revealed a putative interaction of MPV17 through its N-terminal part with MIC60 (**Fig EV.2D**). This interaction of MPV17 with MIC60 was previously demonstrated in mouse cardiomyocytes and could thus be a conserved feature of the protein (Madungwe et al., 2020).

**Figure 3.**
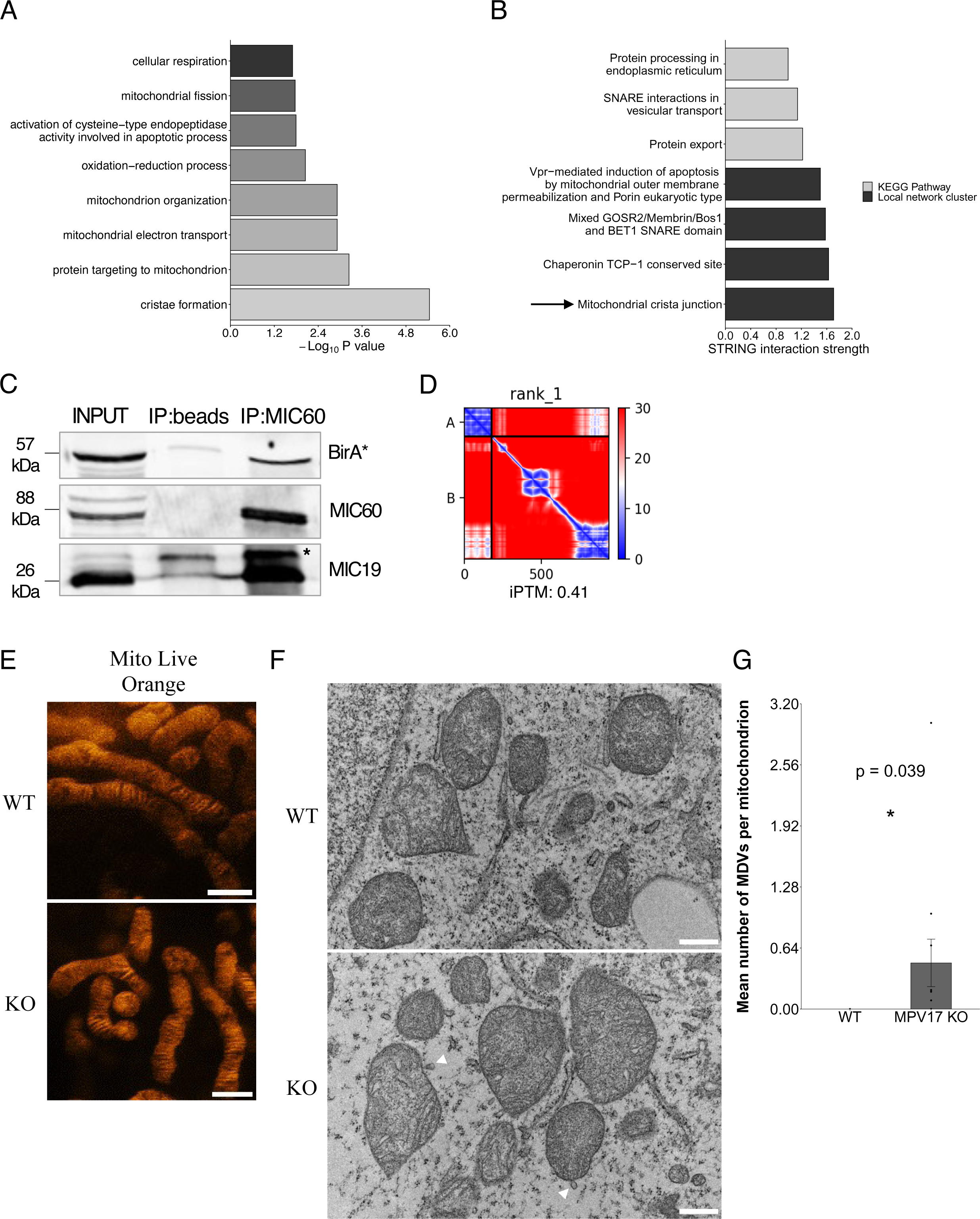
MPV17 interacts with the MICOS complex. (**A**) GO enrichment analysis done on the ≥ 2-fold-enriched proteins in the MPV17-BirA* BioID compared to preOTC-BirA* condition, using the DAVID platform and showing the top enriched BP. (**B**) Interaction network analysis done on the ≥ 2-fold-enriched proteins in the MPV17-BirA* BioID compared to preOTC-BirA* condition, using the STRING platform (von Mering et al., 2003) and showing the top significant terms, either from local network cluster from STRING database or from KEGG database, regarding their STRING interaction strength. (**C**) HEK293T cells were seeded, transfected 24 h later with the MPV17-BirA* construct and harvested 48 h after transfection for protein extraction. 800 µg of proteins were loaded on magnetic beads coupled or not with 5 µg of MIC60 antibody. Proteins were immunoprecipitated for 16 h on a wheel at 4 °C and proteins were resolved by SDS-PAGE. 5 % of the lysate (40 µg) was used as input. BirA*, MIC60 and MIC19 were revealed by Western blot analysis (Representative analysis of three independent experiments). Asterisk corresponding to non-specific band. (**D**) Heatmap obtained with the Predicted Aligned Error (PAE) viewer (Elfmann & Stülke, 2023) for graphic representation of Alphafold multimer prediction (Evans et al., 2022) representing the confidence level of an interaction model between MPV17 and MIC60. The x axis represents a scale based on the number of amino acids of the 2 proteins. The blue squares in the top left and bottom right represent the confidence level of the prediction of the protein alone. The lower left square represents the confidence level of the interaction model between protein B and protein A, and the upper right square represents the confidence level between protein A and protein B. The upper left and lower right corresponds to the self-interaction of the protein. The confidence level is defined by the expected position error of AlphaFold at residue x and is represented by colours ranging from red to blue. An expected position error of up to 30 Å is considered unreliable and is therefore represented in red. The numerical value associated with this confidence level of the interaction model is given by iPTM value, ranging from 0 to 1. (**E**) HEK293T*^MPV17+/+^*and HEK293T*^MPV17-/-^* cells were seeded and stained 24 h after with 100 nM of Live Orange probe for 30 min. Micrographs were acquired 10 min after on an Abberior INFINITY microscope. Representative experiment of three independent biological replicates. Scale bar = 1 µm. (**F**) HEK293T*^MPV17+/+^*and HEK293T*^MPV17-/-^* cells were seeded, fixed 24 h later with 2.5 % electron microscopy-grade glutaraldehyde and prepared for transmission electron microscopy. Micrographs were acquired on a JEOL JEM 1400 transmission electron microscope. Representative experiment of two independent biological replicates. The white arrows depict Mitochondria-Derived Vesicles (MDVs) events that could only be detected in MPV17 KO cells. Scale bar = 500 nm. (**G**) Quantifications of the mean number of MDV events per mitochondrion detected in the TEM micrographs shown in **F**, using Fiji (ImageJ2) software (n = 7 cells for WT and n = 12 cells for KO using two different clones of WT and KO cells). Data are represented as mean ± SEM. * *p < 0.05*.

To investigate MPV17 function in crista formation/maintenance, we took advantage of super-resolution Stimulated Emission Depletion (STED) microscopy and observed the mitochondrial network in MPV17 WT and KO cells (**Fig.3E**). However, no major phenotypical effect of the absence of MPV17 on the mitochondrial ultrastructure was observed. A similar observation was made using Transmission Electron Microscopy (TEM) (**Fig.3F**). Interestingly, MDVs were observed on TEM micrographs (**Fig.3F**) only in MPV17 KO cells (**Fig.3G**), suggesting a higher stress in KO cells compared to WT cells. Importantly, damaged mtDNA can be degraded through MDVs, a process dependent on vesicular transport, as was demonstrated in mouse using proximity labelling of the mtDNA helicase TWINKLE (TWNK) (Sen et al., 2022).

### 4.4 The mtDNA proxisome of MPV17 KO cells highlights an enrichment of DNA repair proteins

In order to get additional clues about the fate of the mtDNA in MPV17 KO cells, we took advantage of the previously published TWNK-APEX2 construct (Sen et al., 2022), to generate a TWNK-BirA* construct acting as a bait for mtDNA proxisome profiling in MPV17 WT and KO cells. We first validated the mitochondrial localization of the construct (**Fig.4A**) as well as its biotinylating activity (**Fig EV.3A**). As the biotinylation was already observable after 6 h of incubation with biotin, we selected this short exposure to avoid putative deleterious effects of the biotinylation on the mtDNA replication and transcription. The label-free quantification of the BioID results highlighted few significant differences in the proxisome of TWNK between the WT and KO cells, with only 122 proteins showing ≥ 1.5-fold-change enrichment, and 95 showing ≤ 0.67-fold-change enrichment (**Fig.4B**, **Dataset EV4**). A Gene Set Enrichment Analysis (GSEA) of those data highlighted the enrichment of the Molecular Function (MF) term “monovalent inorganic cation transmembrane transporter activity” (Normalized Enrichment Score = 1.9), in line with a potential role of MPV17 as channel of the inner membrane (Antonenkov et al., 2015; Sperl & Hagn, 2021).

**Figure 4.**
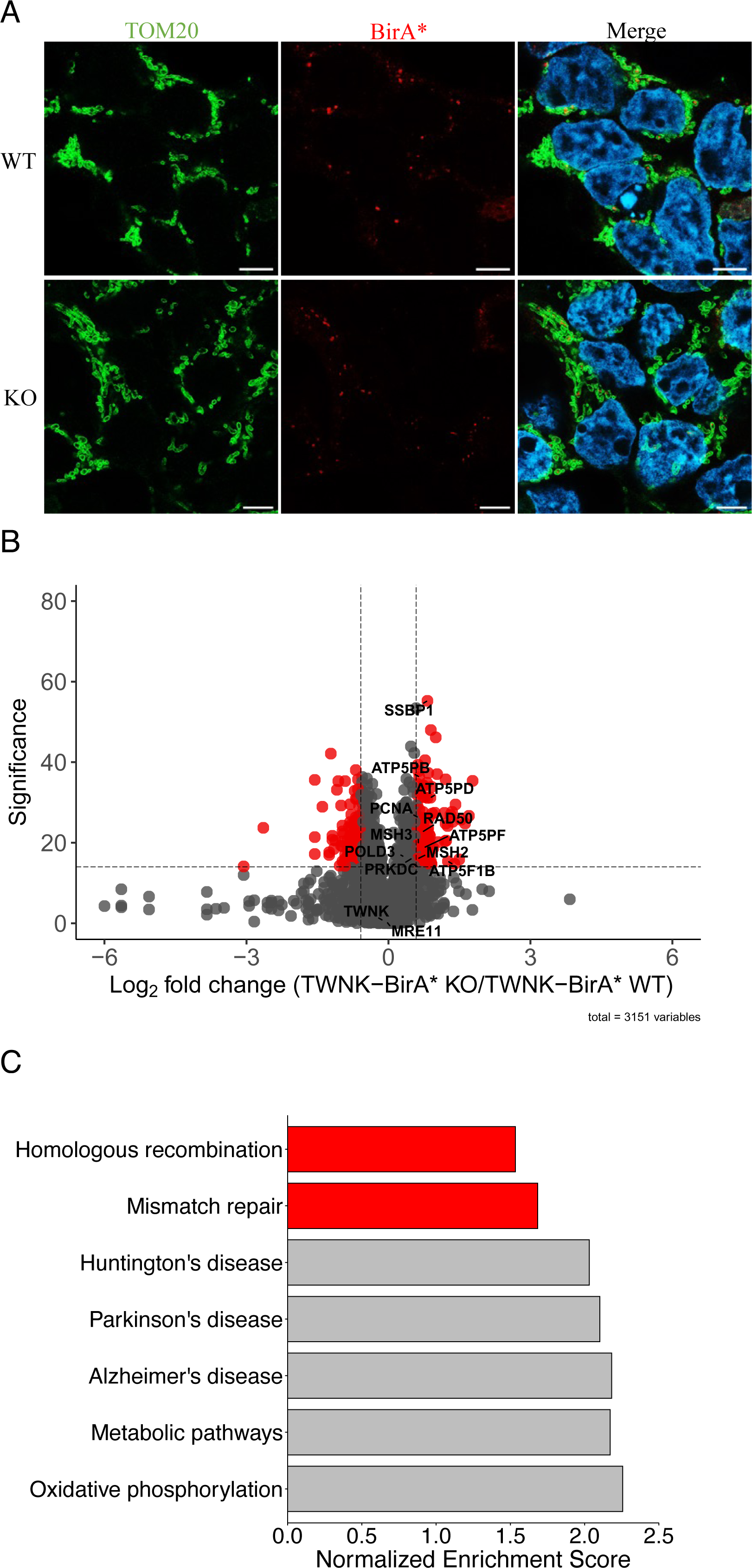
The proxisome of TWNK suggests a potential increased load of mtDNA damages with increased repair. (**A**) HEK293T*^MPV17+/+^*and HEK293T*^MPV17-/-^* cells were seeded and transfected 24 h later with the TWNK-BirA* construct. The cells were fixed 24 h after in 4 % PFA and labelled with anti-TOM20 and anti-BirA* antibodies. Nuclei were stained using DAPI. Micrographs were acquired on a Zeiss LSM900 confocal microscope with an Airyscan detector. Scale bar = 5 µm. Representative experiment of three independent biological replicates. (**B**) HEK293T*^MPV17+/+^*and HEK293T*^MPV17-/-^* cells were seeded and transfected 24 h later with the TWNK-BirA* construct. The cells were incubated 24 h after with 50 µM of biotin for 6 h before harvesting and protein extraction. The maximum amount of proteins (2.3 mg) was loaded on streptavidin-coated beads and the biotinylated proteins were pulled-down for 16 h prior digestion on beads and mass spectrometry analysis, using a nanoLC-ESI-MS/MS tims TOF Pro coupled with an UHPLC nanoElute. The identification and label-free quantifications were done using PEAKS Studio 11 software. A 1% FDR and a single peptide detection threshold were set. Data are represented as a measure of enrichment of proteins in *MPV17* KO cells compared to WT ones (n = 3). Different proteins of interest are displayed on the volcano plot and a 15-significance threshold and 1.5-fold change threshold were set. (**C**) A GSEA was performed, using the KEGG database and the top positively enriched terms with their Normalized Enrichment Score (NES) are displayed.

The GSEA using KEGG database revealed other features differentiating the TWNK proteome of WT and KO cells. First, the “Oxidative phosphorylation” pathway was the top enriched process along with several disease-associated pathways and the general “Metabolic pathways term” (**Fig.4C**), all linked to the enrichment of ATP synthase subunits (**Fig.4B**). This enrichment did not result from a higher abundance of the different subunits, as their abundance was similar in both WT and KO cell fractions (**Fig1.F**), but corresponds to a higher proximity between them and TWNK. The second most enriched list of proteins corresponds to the nuclear process of “Mismatch repair” (illustrated in **Fig EV.3B**) and “Homologous recombination” (**Fig.4C**). Of note, no difference between MPV17 WT and KO cells was observed for the abundance of the different nuclear DDR proteins (**Fig.1F**), suggesting a higher proximity of these proteins with the mtDNA in KO cells. Those results suggest potential damages in the mtDNA of KO cells since nuclear DDR proteins are known to be recruited to mitochondria upon oxidative damages to restore the mtDNA genome integrity (Achanta et al., 2005; Bannwarth et al., 2012; Kalifa et al., 2012; Papeta et al., 2010; Sage et al., 2010), and reinforce the concept that the mtDNA depletion in MPV17 KO cells could be caused by mtDNA damages followed by degradation.

### 4.5 The higher mitophagy observed in MPV17 KO cells is not responsible of the mtDNA depletion

Damaged mtDNA molecules and more generally damaged mitochondria are known to be preferentially degraded through mitophagy (Fu et al., 2020; Shu et al., 2021). Therefore, we assessed mitophagy and also autophagy in MPV17 WT and KO cells, using the Fis1-GFP-mCherry reporter reflecting mitophagy (**Fig.5A**), and LC3B-GFP-mCherry reporter reflecting autophagy (**Fig.5C**). Both reporters are based on the quenching of the GFP signal in acidic compartments and thus the quantification of the mCherry-only puncta is correlated with the mitophagy (for the Fis1-GFP-mCherry reporter) and autophagy (for the LC3B-GFP-mCherry reporter) levels (Allen et al., 2013; Pankiv et al., 2007). We observed a significantly higher level of mitophagy in KO cells compared to the control condition (**Fig.5B**), but no significant increase in the autophagy level (**Fig.5D**). In addition, the mitochondrial biogenesis, assessed by the activity of several transcription factors (NRF1, CREB and YY1) and by the activity of the TFAM promoter, was not significantly different between both cell lines (**Fig.5E**). Interestingly, and as mentioned before, we also observed the presence of MDVs in KO cells (**Fig.3F-G**), witnessing the removal of small (70-150 nm) fragments of mitochondria containing damaged contents (Sugiura et al., 2014). The MDVs observed in MPV17 KO cells might thus contain damaged mtDNA. Indeed, the average area of those MDVs was calculated (0,0128 µm^2^) and could match the size of single mtDNA nucleoids (0.00758 µm^2^ for active nucleoids) (Brüser et al., 2021). However, the absence of MDVs with a double membrane does not support the presence of mtDNA inside (**Fig.3F**). Additionally, blocking either the lysosomal activity, using bafilomycin, or the initiation of autophagy, using ULK1 inhibitor (MRT68921), did not rescue the mtDNA depletion observed in KO cells (**Fig.5F**). Thus, the higher mitophagy and the increased MDV production observed in KO cells is not the primary cause of the mtDNA depletion. In addition, the absence of compensatory biogenesis (**Fig.5E**) and of global autophagy (**Fig.5D**) indicates that the higher degradation of the Fis1-GFP-mCherry reporter could originate from the degradation of mitochondrial fractions, potentially MDVs, rather than whole mitochondria.

**Figure 5.**
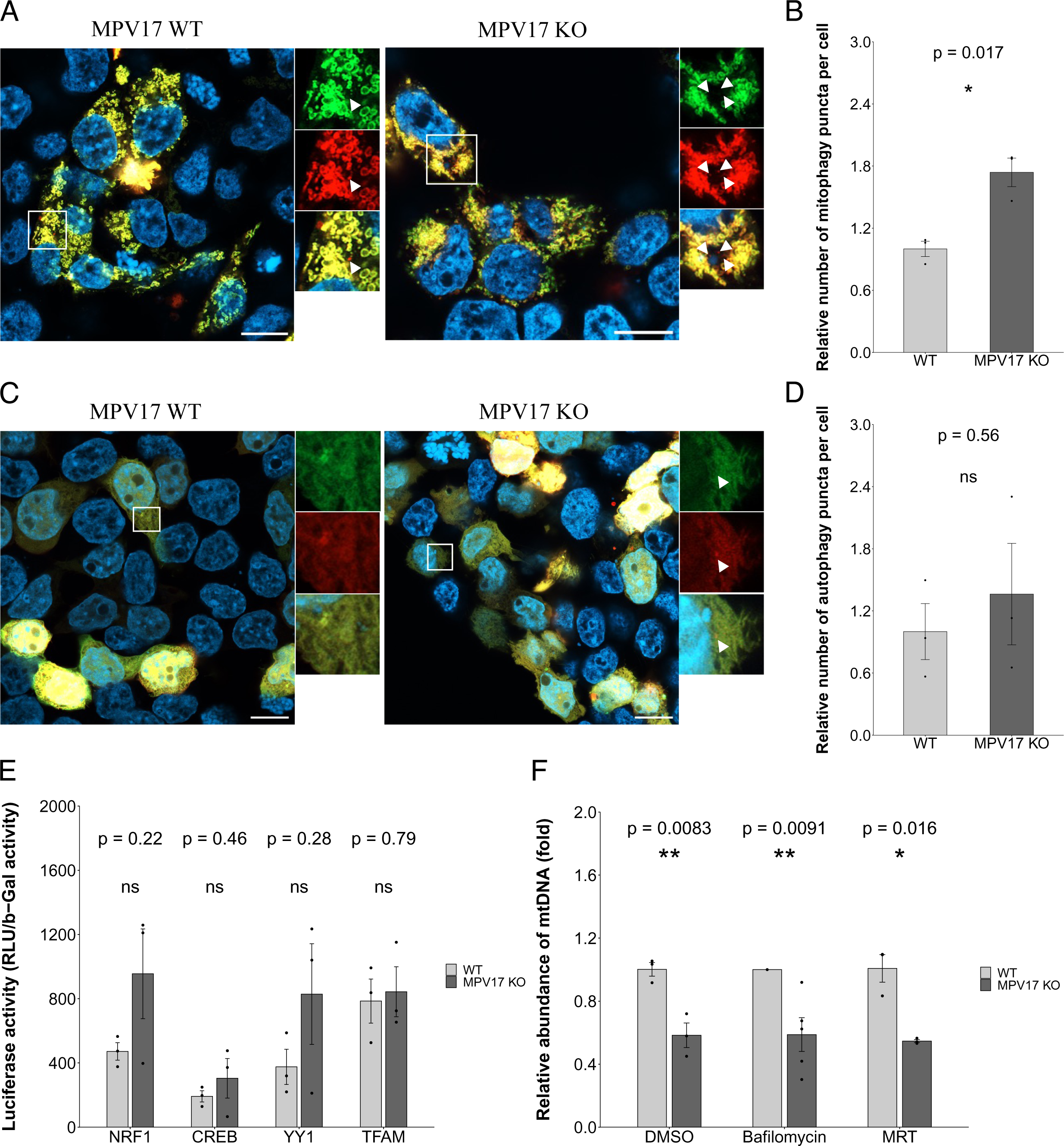
MPV17 KO cells display a higher mitophagy and MDVs number, although unrelated to the mtDNA depletion. (**A-B**) HEK293T*^MPV17+/+^* and HEK293T*^MPV17-/-^* cells were seeded and transfected 24 h later with the Fis1-GFP-mCherry reporter construct (Allen et al., 2013). 48 h after the cells were fixed in 4 % PFA and stained with DAPI. Micrographs were acquired on a Zeiss LSM900 confocal microscope with an Airyscan detector. Scale bar = 10 µm. The mitophagy was quantified by counting the average number of red puncta per cell as shown with the white arrow, using Fiji (ImageJ2) software (**B**). An unpaired t-test was performed and quantifications were done on 20-50 transfected cells per replicate (n = 3 using three different clones of WT and KO cells). Data are represented as mean ± SEM. * *p < 0.05*. (**C-D**) HEK293T*^MPV17+/+^* and HEK293T*^MPV17-/-^* cells were seeded and transfected 24 h after with the LC3B-GFP-mCherry reporter (Pankiv et al., 2007). 48 h after the cells were fixed in 4 % PFA and stained with DAPI. Micrographs were acquired on a Zeiss LSM900 confocal microscope with an Airyscan detector. Scale bar = 10 µm. The autophagy was quantified by counting the average number of red puncta per cell, using Fiji (ImageJ2) software (**D**). An unpaired t-test was performed and quantifications were done on 20-50 transfected cells per replicate (n = 3 using three different clones of WT and KO cells). Data are represented as mean ± SEM. (**E**) HEK293T*^MPV17+/+^* and HEK293T*^MPV17-/-^*cells were seeded and transfected 24 h after with the NRF1, CREB, YY1 or TFAM luciferase reporters in combination with the pCMV-βGal reporter. 48 h after, cells were harvested, lysed in Reporter Lysis Buffer and processed according to manufacturer’s instruction for luciferase and βGal activity measurement. The luciferase reporter’s activity, is expressed as a ratio between the measured relative light units (RLU) and the 405 nm optical density value obtained from βGal measurement. An unpaired t-test was performed for each condition (n = 3 using at least two different clones of WT and KO cells). Data are represented as mean ± SEM. (**F**) HEK293T*^MPV17+/+^* and HEK293T*^MPV17-/-^*cells were seeded and treated 24 h after with either DMSO (vehicle), 50 nM Bafilomycin or 5 µM MRT68921 for 48 h. To be able to observe the depletion phenotype at confluency, 10 nM of ethidium bromide supplemented with 50 µg/mL uridine and 1 mM pyruvate were added for the last 24 h. Cells were then harvested for DNA extraction and the mtDNA abundance was measured using qPCR with primers targeting the mitochondrial genes *ND1* and *ND2* and was normalized using the abundance of the nuclear genes *Beclin* and *TBP*. Data are expressed relative to WT cells and an unpaired t-test was performed for each condition (n = 3 for DMSO and MRT and n = 5 for Bafilomycin condition using at least two different clones of WT and KO cells). Data are represented as mean ± SEM. * *p < 0.05*, ** *p < 0.01*.

### 4.6 In silico analyses of MPV17 structure and partners support a calcium channel function

To further get insights in MPV17 molecular function, we used the Alphafold multimer prediction tool to test 2-by-2 dimerization of MPV17 with candidates enriched in the MPV17-BioID dataset. These candidates were selected among the ≥ 2-fold-change enriched proteins on the basis of their biological function. More precisely, selected proteins had a potential function in cristae organization; a putative association with the peroxisome, already described for its homolog MPV17-like (Iida et al., 2006); an association with the mitochondrial permeability, previously described in mouse (Madungwe et al., 2020) and suggested by the STRING analysis (**Fig.2B**); or an association with mitochondrial quality control systems (**Table EV4**). The iPTM value of each interaction was calculated and the arbitrary threshold corresponding to the iPTM value of MIC60 (0.41) was considered to sort the most interesting candidates (**Table EV4**). Among those, the CyclophilinD (CypD), the main regulator of the mPTP (Hurst et al., 2017), drew our attention, because an interaction with CypD would support the proposed role of MPV17 as a cation channel embedded in the inner membrane (Antonenkov et al., 2015). In addition, an enrichment of cation channel activity molecular function (MF) was observed in the proxisome of TWNK in MPV17 KO cells (**Fig EV.3B**) which suggests a compensatory phenotype. The heatmap-based representation of the interaction between MPV17 and the CypD suggests a likely interaction between both proteins (**Fig.6A**), which was experimentally confirmed by co-immunoprecipitation (**Fig.6B**). The 3D modeling of the interaction further suggests that the interaction occurs at the C-terminal of MPV17 with the S170 residue interacting closely to CypD active site, represented by the R55 residue (**Fig.6C**). Additional *in silico* analysis further supports a channel conformation with many conserved residues located inside the channel structure (**Fig.6D**). The modeling of the channel depicts a hollow cylinder, featuring a maximal opening of 12 Å (1.2 nanometers) situated at the N-terminal region. Progressing towards the C-terminus, the cylinder diameter exhibits a gradual reduction. Notably, a narrow bottleneck with a diameter of merely 3 Å was identified in the plane intersecting horizontally at the 104^th^, 54^th^, and 169^th^ amino acid residues (**Fig.6E**). In addition, predicted impacts on the protein structure of some mutations found in patients revealed a major atomic deviation of the S170F variant of the protein, related to a severe phenotype in patients (El-Hattab et al., 2010; El-Hattab et al., 2018), with a Root Mean Square Deviation (RMSD) value of 3.44 (**Fig. 6F**). This structural deviation fits with a predicted structural role of the S170 residue (**Fig.6D**). At the opposite, the P98L mutation displays almost no modification of the structure (**Fig.6F**, RMSD of 0.18) and is clinically associated with a milder phenotype (Blakely et al., 2012; El-Hattab et al., 2018). Of note, the deviation observed for the S170F mutation occurs in the C-terminal part of the protein (**Fig.6F**) and may thus be associated with a gain or loss of association with the CypD. Altogether these results suggest an interaction of MPV17 with CypD, highlighting a putative functional association of MPV17 with the mPTP.

**Figure 6.**
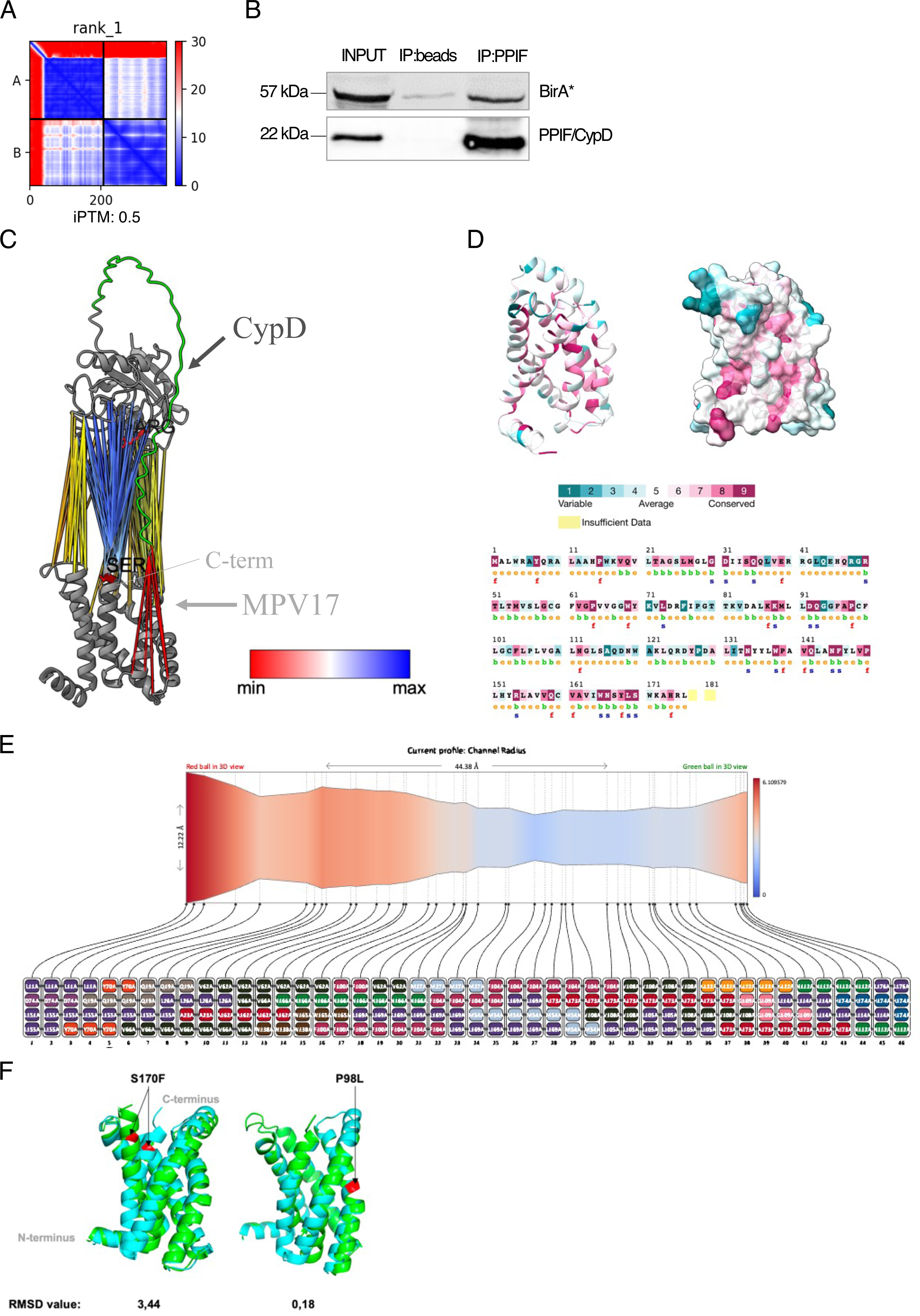
Alphafold-based prediction of MPV17 structure and interacting partners supports a channel activity of the protein. (**A**) PAE heatmap obtained from Alphafold multimer prediction (Evans et al., 2022) representing the confidence level of an interaction model between MPV17 and CypD. (**B**) HEK293T cells were seeded, transfected 24 h later with the MPV17-BirA* construct and harvested 48 h after transfection for protein extraction. An amount corresponding to 600 µg of proteins was loaded on magnetic beads coupled or not with 5 µg of PPIF/CypD antibody. Proteins were immunoprecipitated for 16 h on a wheel at 4 °C and proteins were resolved by SDS-PAGE. 5% of the lysate (30 µg) was used as input. BirA* and PPIF/CypD were revealed by Western blot analysis (n = 3). (**C**) Representation of the predicted dimerization model of MPV17 (light grey) with CypD (dark grey), using ChimeraX software (Meng et al., 2023) and based on the prediction produced with the Alphafold multimer prediction tool. The colored links depict the predicted interaction between amino acids with a confidence level ranging from low (red) to high (blue), showing an interaction of CypD with the C-terminal part of MPV17. The site-active CypD ARG55 residue and the conserved SER170 residue of MPV17 are highlighted. (**D**) Representation of conserved amino acids in MPV17 following sequence alignment using the ConSurf server (Ashkenazy et al., 2016) and using 150 input sequences. The letters below the conserved amino acids describe an exposed amino acid (e), a buried residue (b), a residue predicted to be associated with a functional site of the protein due to its high conservation and exposure (f), and a residue predicted to be important to the structure of the protein due to its high conservation and burial (s). (**E**) Graphical representation of the MPV17 channel showing length and diameter, done using the ChExVis tool (Masood et al., 2015). The colors are based on the radius of the channel, ranging from blue for a small radius to red for a larger radius. The amino acid residues forming each horizontal section of the protein are shown below. (**F**) Visualization of the impact of the (from left to right) S170F and P98L mutations on MPV17 predicted structure. The WT is shown in green, and the mutated protein is shown in blue. The mutated amino acids are colored in red. Protein alignment and Root Mean Square Deviation (RMSD) calculations were performed using PyMOL (v2.5). High (close to 4) RMSD values are considered as significant conformation changes.

### 4.7 Involvement of MPV17 in the mitochondrial calcium regulation through mPTP association

Since the mPTP generates fast calcium efflux from mitochondria, we assessed the mitochondrial calcium level in MPV17 WT and KO cells. Using the Rhod2AM calcium probe and its property to accumulate inside mitochondria (Zhu et al., 2000), we measured the mitochondrial calcium level in both cell lines, using Rhodamine 123 as a mitochondrial marker (**Fig.7A**). Quantification of the mitochondrial Rhod-2 AM signal highlighted a higher calcium level in KO cells, suggesting a potential role of MPV17 as a calcium efflux channel (**Fig.7B**). We reasoned that in the presence of higher mitochondrial calcium level, the opening of the mPTP would be more easily triggered following intracellular calcium rise, as for example using ionomycin treatment which rises intracellular calcium level, then the mitochondrial calcium level to finally open the mPTP (Fernandez-Sanz et al., 2019; Morgan & Jacob, 1994; Peng & Greenamyre, 1998). To assess WT and KO cells response to ionomycin-induced mPTP opening, we used the mito-GCaMP5G, a mitochondrially targeted calcium reporter construct (Kwon et al., 2016). In response to ionomycin, we observed a rise in fluorescence, corresponding to mitochondrial calcium influx, rapidly followed by a fluorescence drop related to mPTP opening (**Fig.7C**). Timelapse recording revealed a high intercellular variability in terms of reporter expression and calcium influx and efflux kinetics (**EV movies 1-4**). However, comparison of the mPTP opening sensitivity between the different experimental conditions revealed a higher resistance of mPTP opening in KO cells compared to WT ones, but similar to the one observed for WT cells treated with cyclosporin A (CsA), a CypD inhibitor reducing mPTP opening sensitivity (Bernardi et al., 2023; Tanveer et al., 1996) (**Fig.7D**). Interestingly, CsA could not increase the mPTP resistance of KO cells, suggesting that MPV17 acts downstream of CypD and could be responsible for the CsA-sensitivity of the mPTP opening in those cells.

**Figure 7.**
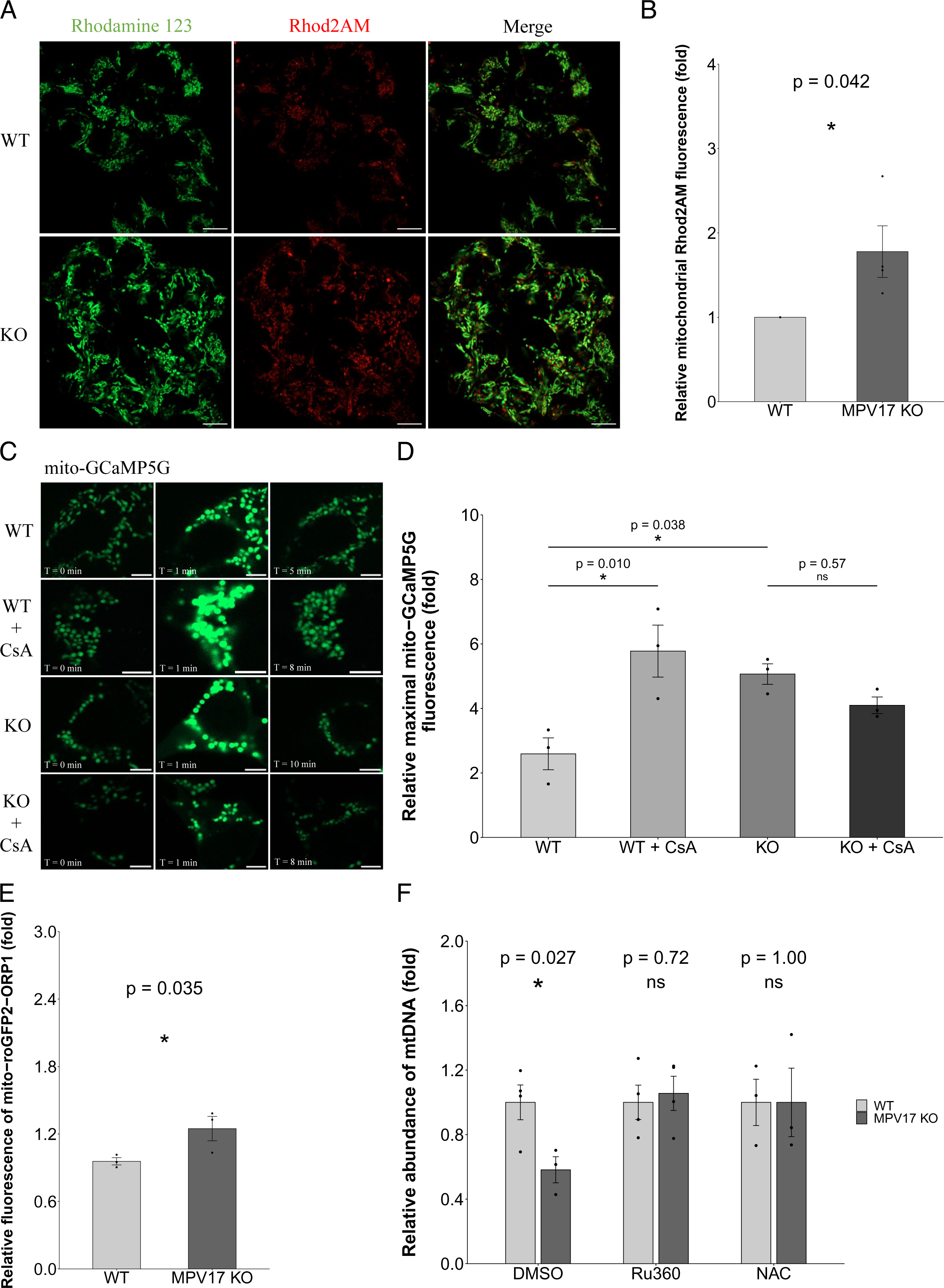
MPV17 involvement in mitochondrial calcium homeostasis and association with mPTP function. (**A-B**) HEK293T*^MPV17+/+^* and HEK293T*^MPV17-/-^* cells were seeded and stained 6 h after with 6 µM of Rhod2AM probe for 1 h at 37 °C priori medium refreshing and incubation for 18 h. Cells were then co-stained with 10 µg/mL of Rhodamine 123 for 10 min prior imaging using a Zeiss LSM900 confocal microscope. Scale bar = 10 µm. Quantification of the Rhod2AM fluorescence colocalizing with Rhodamine 123 signal (mitochondrial mask) was performed using Fiji (ImageJ2) software (**B**). An unpaired t-test was performed and quantifications were done on 75-100 cells per replicate (n = 4, using two different clonal populations). Data are represented as mean ± SEM. * *p < 0.05*. (**C**) HEK293T*^MPV17+/+^*and HEK293T*^MPV17-/-^* cells were seeded and transfected 24 h after with the mito-GCaMP5G reporter (Kwon et al., 2016). 48 h after, the cells were treated with 500 nM ionomycin in combination or not with 5 µM cyclosporin A (CsA) and the fluorescence intensity of the construct was measured every min for 30 min following treatment. Micrographs were acquired on a Zeiss LSM900 confocal microscope with an Airyscan detector. Scale bar = 5 µm. The micrographs show representative examples of basal fluorescence (left panels, time (T) = 0 min), maximal fluorescence following treatment (middle panels) and residual fluorescence following mPTP opening and calcium efflux (right panels), with their corresponding timing. (**D**) The maximal fluorescence intensity relative to T = 0 min of each cell was calculated using Fiji and corresponds to the maximal calcium threshold required to open the mPTP in the different conditions. An ANOVA followed by a TukeyHSD post-hoc test was performed and quantifications were done on 3-4 cells per replicate (n = 3 using two different clonal populations of WT and KO cells). Data are represented as mean ± SEM. (**E**) HEK293T*^MPV17+/+^* and HEK293T*^MPV17-/-^* cells were seeded and transfected 24 h after with the mito-roGFP2-ORP1 reporter (Werley et al., 2020). 48 h after, the fluorescence intensity of the construct was measured using a Zeiss LSM900 confocal microscope with an Airyscan detector. The fluorescence of the construct excited at 405 nm was normalized by the fluorescence following excitation at 488 nm and the normalized fluorescence was expressed relatively to the control. A paired t-test was performed and quantifications were done on 75-100 cells per replicate (n = 3, using two different clonal populations). Data are represented as mean ± SEM. * *p < 0.05*. (**F**) HEK293T*^MPV17+/+^*and HEK293T*^MPV17-/-^* cells were seeded and treated 24 h after with either DMSO (vehicle), 5 µM Ru360 or 4 mM NAC for 72 h. To be able to observe the depletion phenotype at confluency, 10 nM of ethidium bromide supplemented with 50 µg/mL uridine and 1 mM pyruvate were added for the last 24 h. Cells were then harvested for DNA extraction and the mtDNA abundance was measured using qPCR with primers targeting the mitochondrial genes *ND1* and *ND2* and was normalized using the abundance of the nuclear genes *Beclin* and *TBP*. Data are expressed relative to WT cells and an unpaired t-test was performed for each condition (n = 3 for DMSO KO and n = 4 for DMSO WT and Ru360 and n = 3 for NAC condition using at least two different clonal populations for WT and KO cells). Data are represented as mean ± SEM. * *p < 0.05*.

Thus, in HEK293T cells, MPV17 acts as a calcium efflux channel through the IMM, which is associated with the mPTP and would be regulated by CypD. To link this function to the mtDNA depletion observed in the KO cell lines, we investigated the calcium-induced ROS production, as high calcium levels enhance the activity of Krebs cycle dehydrogenases and increase electron leakage leading to ROS production (Brookes et al., 2004; Denton et al., 1978; McCormack & Denton, 1979). Moreover, ROS can damage mtDNA, leading to its repair or degradation (reviewed in Rong et al., 2021). Using the H_2_O_2_ sensor roGFP2-ORP1 (Werley et al., 2020) we noticed higher ROS levels in KO cells when compared to WT ones (**Fig.7E**). In parallel we disclosed that the inhibition of calcium entry, using the MCU inhibitor Ru360, or the treatment with the N-acetyl-cysteine (NAC) antioxidant rescues the mtDNA depletion observed in KO cells (**Fig.7F**). These results support a link between higher mitochondrial calcium levels and a ROS-dependent mtDNA depletion in MPV17 KO cells, which we summarized in **Figure 8**.

**Figure 8.**
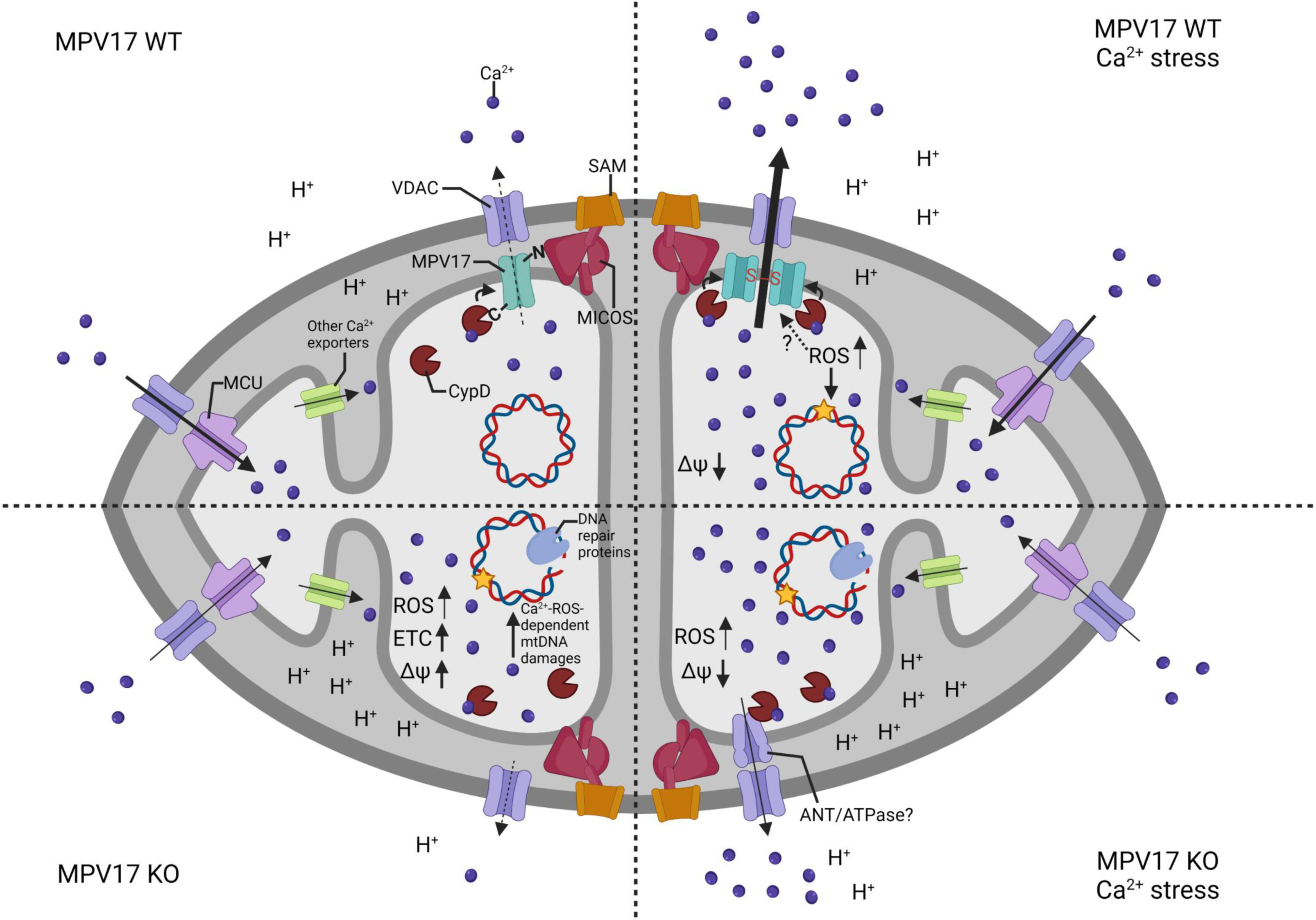
Model for MPV17 as a calcium efflux channel of the IMM associated with the mPTP. The model for MPV17 function presented in this study proposes a role of calcium efflux channel for the protein which would be regulated by CypD under calcium stress conditions, promoting its calcium-dependent opening involved in mitochondrial permeability transition. Moreover, the protein was previously described to oligomerize upon oxidative stress and could also be involved in the ROS-sensitive mPTP opening (Corra et al., 2023; Sperl & Hagn, 2021). In the absence of the protein, the calcium builds up in the mitochondria, leads to increased electron transfer, and subsequent higher membrane potential, increasing ROS production through electron leaking. The constant presence of oxidative damages on mtDNA leads to the recruitment of DNA damage response proteins for repair and can lead to degradation of unrepaired molecules, responsible for the mtDNA depletion observed in KO cells. Upon calcium stress, the cells would present a higher resistance to CypD-mediated mPTP opening which would open alternatively in a CypD-independent manner. Besides, MPV17 is also associated with the MICOS and directly interacts with MIC60 at crista junctions, close to active mtDNA nucleoids. We propose that MPV17 interacts with MIC60 through its N-terminal domain, which would be in the intermembrane space, and with CypD through its C-terminal domain which would thus be in the matrix side. The figure was generated on Biorender (https://app.biorender.com/).

## 5. DISCUSSION

In this study, we used a human cell line as a model but the initial characterization of the HEK293T*^MPV17-/-^* cell lines revealed only a mild phenotype with few alterations of the proteome. However, a mtDNA depletion was nonetheless observed, exclusively in proliferative cells, and was rescued with both tagged and untagged forms of human MPV17. According to the study of Dalla Rosa and colleagues, the mtDNA depletion occurs mainly in quiescent cells and is linked to the depletion of the mitochondrial deoxynucleotide pool (Dalla Rosa et al., 2016). Those observations were made in *MPV17*^-/-^ mouse and in *MPV17*-mutated patient fibroblasts. Unexpectedly, we were unable to observe any effect of a deoxynucleotide supplementation on the recovery of mtDNA following depletion (**Fig.1C**). In addition, we observed the mtDNA depletion only in proliferative cells and the inhibition of the mtDNA replication with ethidium bromide did not equalize the mtDNA levels between WT and KO cells. A similar observation was made in patient-derived fibroblasts displaying POLG mutations which showed altered or slower mtDNA replication (Stewart et al., 2011), as proposed to occur in MPV17 KO/mutant models. In our cellular model, the deficit of replication affected only the recovery of the mtDNA after depletion. In addition, when we treated cells at confluency, presenting comparable mtDNA abundance, with ethidium bromide described to inhibit the mtDNA replication, we phenocopied the depletion observed in proliferative cells (**Fig.1D**). We thus propose that the mtDNA depletion from proliferative cells where the replication capacity of the mtDNA is maximal, and in cells treated with ethidium bromide, is due to a higher degradation rate, rather than to a deficit in biosynthesis. To explain the difference in mtDNA levels between the different models of MPV17 disease, the mtDNA degradation hypothesis can still hold. Indeed, it was demonstrated that, upon mtDNA insults, such as oxidative damages, the rate of mtDNA degradation and the rate of mtDNA recovery depend on the cell type, explaining the multiple different phenotypes associated with MPV17 invalidation or mutation (Friese et al., 2019). The first characterization of the KO cell lines was quite unexpected since a similar model using a shRNA-based constitutive knock-down of MPV17 in HEK293T cells displayed results different from what we observed. Indeed, Jacinto and colleagues showed that MPV17 silenced cells do not display mtDNA depletion, while presenting a quiescent metabolic profile (Jacinto et al., 2021), contrarily to the higher membrane potential we observed in the MPV17 KO cells. However, it is not clear at which stage of confluence they performed their mtDNA measurement in their study, as mtDNA level assessment in confluent cells would be in agreement with our results. Most importantly, our group, using a similar shRNA approach for MPV17 invalidation, confirmed mtDNA depletion in MPV17-silenced cells and observed a phenotype of reduced cell proliferation in multiple human cancer cell lines. However, we failed to associate this phenotype to MPV17 as it could not be rescued by the overexpression of the protein (Canonne et al., 2020). This stresses the absolute necessity to rescue phenotypes, specifically while performing lentiviral-based invalidation of proteins, to avoid off-target effects. On the contrary, the mtDNA depletion phenotype observed here in MPV17 KO cells was rescued by either tagged or untagged MPV17.

The degradation hypothesis of mtDNA is further supported by the higher ROS level observed in KO cells (**Fig.7E**). This parallels an increased recruitment of DDR proteins as well as mismatch repair proteins in the vicinity of the mtDNA of MPV17 KO cells, supporting the hypothesis of constitutive damages in the absence of MPV17. Indeed, DDR and mismatch repair proteins were already reported to be recruited upon mtDNA damages and to be involved in mtDNA integrity maintenance (Achanta et al., 2005; Bannwarth et al., 2012; Kalifa et al., 2012; Papeta et al., 2010; Rashid et al., 2019; Sage et al., 2010). Interestingly, we observed, in MPV17 proxisome, an enrichment in the DDR protein DNA-PKcs whose mitochondrial localization was confirmed by immunofluorescence (**Fig EV.1C**) and interestingly, both MPV17 and DNA-PKcs are genetically associated in an Adriamycin (ADR)-induced nephropathy mouse model (Papeta et al., 2010).

It was previously demonstrated that oxidative damages on the mtDNA as well as point mutations are selectively removed by MDVs and autophagy-dependent mechanism in mouse fibroblasts (Sen et al., 2022). In the present study, we excluded that the mtDNA degradation would occur through this mechanism, since we did not observe a higher number of cytosolic DNA nor late endosomes containing mtDNA in KO cells, despite observing a higher number of MDVs compared to WT cells. In addition, even though a higher mitophagy rate was observed in KO cells, the mtDNA depletion was not rescued by the inhibition of autophagy or lysosomal activity. Thus, the mechanisms involving a selective removal of oxidized mtDNA by a VPS35-ATAD3-TWNK axis or by ATAD3B-dependent mitophagy (Shu et al., 2021), do not explain the mtDNA depletion in MPV17 KO cells, even though ATAD3B was found highly enriched in MPV17 proxisome (**Dataset EV2**). The higher production of MDVs is correlated with increased mitophagy and is consistent with higher ROS levels, which can oxidize all kinds of biomolecules, such as nucleic acids, proteins, and lipids. Indeed, MDVs are typically formed in response to oxidation and drive the degradation of oxidized proteins and lipids through lysosome targeting (Quiles & Gustafsson, 2020; Sugiura et al., 2014; Todkar et al., 2021). The MDVs observed here could thus contain proteins or lipids but not mtDNA, which would rather be degraded inside the mitochondria. Indeed, linearized mtDNA can be degraded by endogenous nucleases or by the replication machinery itself (Nissanka et al., 2018; Peeva et al., 2018; Zhao, 2019).

The observed interaction of MPV17 with the MICOS complex was reported in a mouse model of cardiomyocyte (Madungwe et al., 2020) and is confirmed in this study in human cells. However, despite a major impact of MPV17 depletion on the mitochondrial ultrastructure in several models such as yeast (Dallabona et al., 2010), mouse (Luna-Sanchez et al., 2020) and zebrafish (Bian et al., 2021), our results suggested that the MPV17-MICOS association might not be relevant in human HEK293T cells.

By using the combination of MPV17-BioID and *in silico* approach, we disclosed a list of partners that were validated, for some. Among those, the interaction of MPV17 with CypD was strongly supported in human HEK293T cells by i) the BioID results, ii) *in silico* prediction and iii) co-immunoprecipitation experiments, in line with the interaction previously described in mouse (Madungwe et al., 2020). In addition, the *in silico* modeling of MPV17 depicts the protein as a channel where the conserved residues buried inside the protein are the constituent of a channel structure. Moreover, we observed a correlation between the predicted impact of a MPV17 mutation on its structure with the severity of the clinical phenotype associated with the mutation, with the most impacting mutation affecting MPV17 C-terminal, potentially involved in CypD interaction.

The predicted channel structure of the protein, previously proposed to act as a cation channel (Antonenkov et al., 2015), together with its potentially functional interaction with the CypD, suggest a relevant link between MPV17 and the mPTP. Moreover, the higher calcium content observed in MPV17 KO cells is in line with a putative function of MPV17 as an efflux calcium channel embedded in the IMM. A higher mitochondrial calcium content would also enhance the activity of calcium-sensitive dehydrogenases of the tricarboxylic acid cycle (Brookes et al., 2004; Tanwar et al., 2021), henceforth causing an increase in the electron transport chain (ETC) activity with a resulting higher membrane potential, as observed in KO cells. Under normal condition, the build-up of mitochondrial calcium triggers the opening of the mPTP which enables the efflux of the accumulated calcium and leads to membrane depolarization (Hurst et al., 2017). The higher mitochondrial calcium levels, together with the higher resistance of MPV17 KO cells to mPTP opening upon ionomycin-induced calcium stress, support a role of MPV17 as a member or partner of the mPTP. Interestingly, treatment with CsA did not increase further the resistance of KO cells to mPTP opening, suggesting that MPV17 is a CypD-regulated calcium channel, member of the mPTP. Besides, both the ATP synthase and the different ANT proteins are known calcium sensitive mPTP members and could mediate the opening of the mPTP in MPV17 KO cells (Giorgio et al., 2013; Karch et al., 2019; Mnatsakanyan et al., 2019; Urbani et al., 2019). Interestingly, in the study of Karch and colleagues, the removal of all ANTs isoforms was not sufficient to abolish mPTP opening and required the additional removal of CypD, suggesting that another CypD-regulated IMM channel exists and could correspond to the ATP synthase and/or MPV17 (Karch et al., 2019). In addition, the proximity of ATP synthase subunits to the mtDNA proxisome in KO cells could result from a compensatory process related to the loss of a calcium-sensitive calcium efflux channel. Indeed, ATP synthase monomers and dimers are ensuring calcium efflux and are part of the mPTP (Giorgio et al., 2013; Mnatsakanyan et al., 2019) and could be recruited to the crista junction of KO cells, close to mtDNA, to promote the maintenance of calcium homeostasis.

In accordance with the literature (Brookes et al., 2004), we propose a link between the higher mitochondrial calcium level with the higher ROS level in MPV17 KO cells, which might then be responsible for mtDNA constant insults, and repair by DDR proteins recruited to mitochondria. This model is supported by the fact that, by preventing the accumulation of calcium inside mitochondria, by MCU inhibition, the mtDNA depletion phenotype observed in KO cells was rescued. Interestingly, the study of Madungwe and colleagues proposed a model where MPV17 is associated with the mPTP, but would act as a negative regulator of the channel under ischemia-reperfusion. This would lead to a complete opening of the mPTP with lesser calcium retention capacity in the MPV17 mutated mouse heart (Madungwe et al., 2020). This apparent contradiction can be explained by the possible different level and type of stress generated in this study compared to the calcium stress generated here, using ionomycin. The discrepancy between these observations could also result from tissue and/or species specificity. Finally, the absence of MPV17 might also be somehow compensated by other proteins such as the ATP synthase complex, resulting in a milder phenotype compared to models using mutated forms of the protein that are still expressed but which could act as dominant negative. This would explain the restricted pathological phenotype observed for *MPV17^-/-^*mouse model, when compared to MPV17-mutated human patients presenting heavy-burden symptoms leading to fatal issues (El-Hattab et al., 2010; Spinazzola et al., 2006).

In addition, modeling the interaction of MPV17 with MIC60 and CypD leads to propose a putative orientation of the protein, while the crystallographic structure and topology of the protein is extremely difficult to obtain and where only limited biophysical studies have been conducted for MPV17 structure unraveling (Sperl & Hagn, 2021). Indeed, based on the MPV17 partners validated by immunoprecipitation, we suggest that the N-terminal domain is located in the intermembrane space (IMS) and the C-terminal domain in the matrix, with the five transmembrane domains spanning the inner mitochondrial membrane. Indeed, MIC60 which interacts with the N-terminal domain, is an IMS protein whereas the CypD, interacting with the C-terminal end, is a matricial protein. This prediction will be useful for further *in cellulo* molecular characterization of the channel function.

In conclusion, by characterizing the HEK293T MPV17 KO cell line, we highlight new features of MPV17 structure and function, disclosing a high molecular complex association with the MICOS and mPTP complexes, and a channel function, contributing to mitochondrial calcium homeostasis (**Fig.8**).

## Supporting information

Supplementary datasets corresponding to Dataset EV1-4

## 6. ACKNOWLEDGEMENT

We thank Dr. Damien Coupeau (UNamur, Belgium) for his great help in the cloning strategies. We thank the Dr. Pr. David Pla Martin (University of Düsseldorf, Germany) for providing us with the pFis1-GFP-mCherry and pLC3B-GFP-mCherry plasmids. We also thank the MorphIm platform (UNamur, Belgium) for the use of confocal and STED microscopes. We finally thank Florence Manero and the SCiAM platform (Université d’Angers, France) for the TEM preparation of samples and for the acquisition of the micrographs.

## 7. AUTHORS CONTRIBUTION

Author Contributions: Conceptualization, S.M., M.C. and P.R.; methodology, S.M. and I.A.; software, S.M., I.A. and M.D.; formal analysis, S.M., L.M., A.C. and I.A.; investigation, S.M., L.M. and I.A.; resources, S.M. and M.D.; data curation, S.M., A.C.; writing—original draft preparation, S.M.; writing—review and editing, T.A., G.L. and P.R.; visualization, S.M.; supervision, T.A. and P.R.; project administration, S.M. and P.R. All authors have read and agreed to the published version of the manuscript.

Funding acquisition: This research received no external funding. S.M. is a research fellow of the Fonds National de la Recherche Scientifique (FNRS), Belgium.

## TABLES AND THEIR LEGENDS

**Table EV1.** Detection of MPV17 in MPV17 WT and KO cell lines using mass spectrometry.

| Conditions | MPV17 spectral count |
| --- | --- |
| Clone 13 MPV17 WT | 7 |
| Clone 5 MPV17 WT | 5 |
| Clone 16 MPV17 WT | 5 |
| Clone 52 MPV17 KO | 0 |
| Clone 44 MPV17 KO | 0 |
| Clone 3 MPV17 KO | 0 |
| Clone 8 MPV17 KO | 0 |

**Table EV2.**
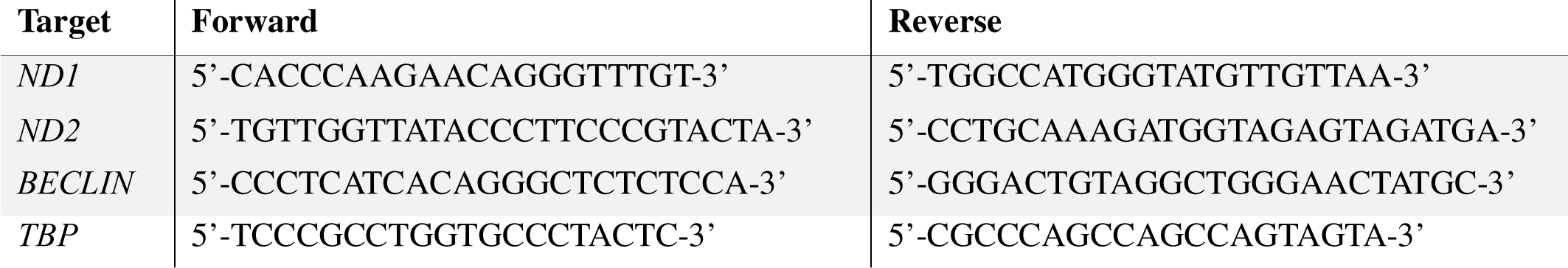
List of used primers.

**Table EV3.** Co-immunoprecipitation of endogenous MPV17 with MIC60 and detection by mass spectrometry.

| Conditions | MIC60 spectral count | MPV17 spectral count |
| --- | --- | --- |
| IP:beads n1 | 22 | 0 |
| IP:MIC60 n1 | 2431 | 1 |
| IP:beads n2 | 112 | 0 |
| IP:MIC60 n2 | 3381 | 2 |

**Table EV4.** MPV17 iPTM interaction scores with selected proteins enriched in the MPV17 BioID.

| Protein_Protein interaction | iPTM |
| --- | --- |
| MPV17_ACBD5 | 0.57 |
| MPV17_IMMP2L | 0.503 |
| MPV17_CYPD | 0.5 |
| MPV17_MIC60 | 0.41 |
| MPV17_YME1L1 | 0.38 |
| MPV17_TFAM | 0.35 |
| MPV17_ATAD3B | 0.339 |
| MPV17_ATAD3A | 0.286 |
| MPV17_OMA1 | 0.24 |
| MPV17_BAX | 0.224 |
| MPV17_BAK | 0.222 |
| MPV17_EXOG | 0.209 |
| MPV17_MPV17 | 0.207 |
| MPV17_MPC2 | 0.2 |
| MPV17_MGARP | 0.19 |
| MPV17_MIC25 | 0.16 |
| MPV17_DIABLO | 0.16 |
| MPV17_MIC26 | 0.14 |

## EXPANDED VIEW FIGURE LEGENDS

**Expanded View Figure 1.**
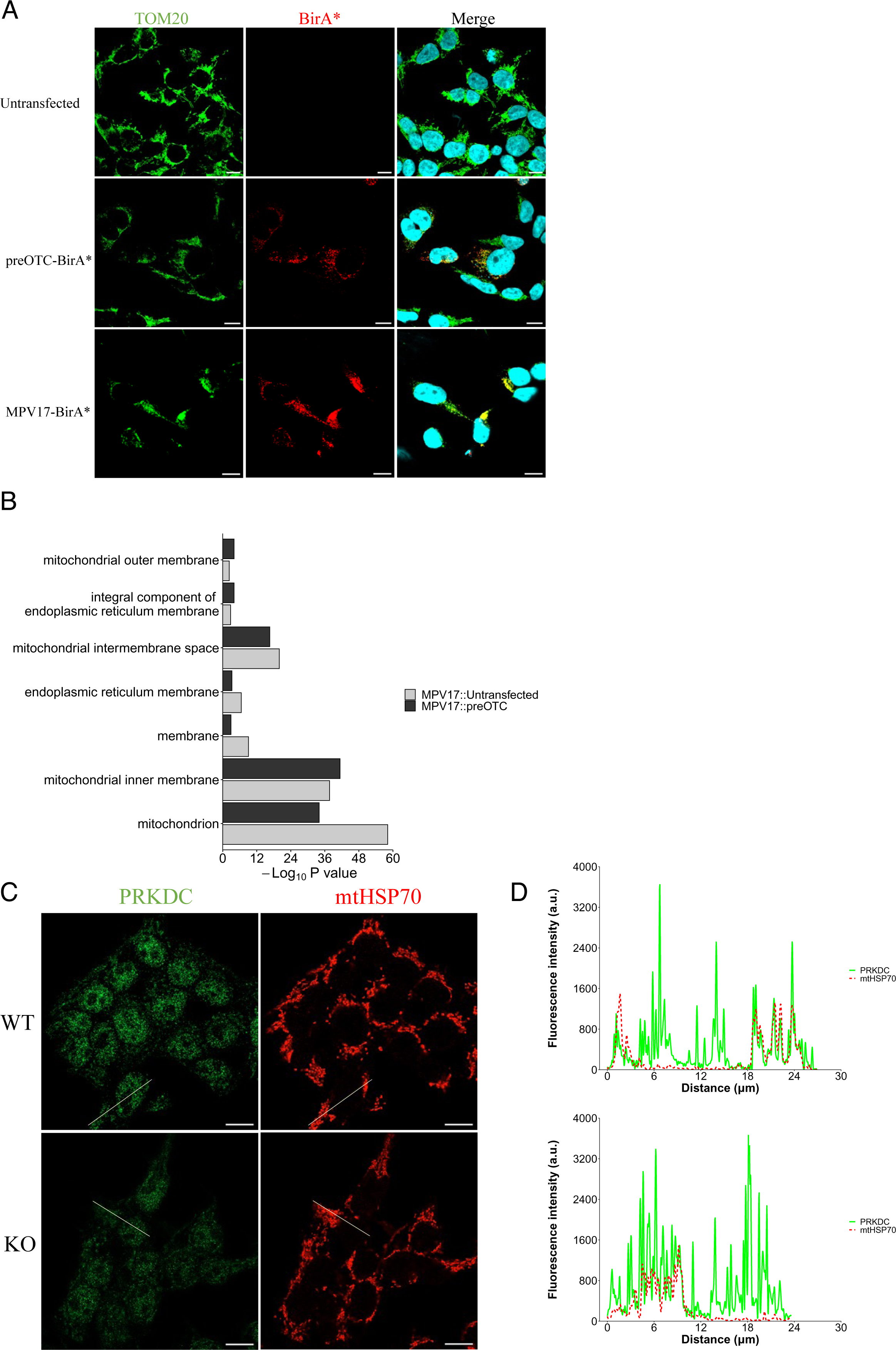
Preliminary analyses of MPV17-BirA* BioID. (**A**) HEK293T cells were seeded and transfected or not 24 h later with either MPV17-BirA* or preOTC-BirA* constructs. The cells were incubated 24 h after with 50 µM of biotin for 18 h before being fixed in 4 % PFA and labelled with anti-TOM20 and anti-BirA* antibodies. Nuclei were stained using DAPI. Micrographs were acquired on a Leica TCS SP5 confocal microscope. Scale bar = 10 µm. Representative experiment of three independent biological replicates. (**B**) Comparison of the GO enrichment analysis done on the ≥ 2-fold-enriched proteins in the MPV17-BirA* BioID compared to either preOTC or untransfected condition. The analysis was done using DAVID platform (Dennis et al., 2003), and shows the top enriched Cellular Component (CC) GO terms. (**C-D**) HEK293T*^MPV17+/+^* and HEK293T*^MPV17-/-^* cells were seeded, fixed 24 h later with 4 % PFA and labelled with anti-DNA-PKcs/PRKDC and anti-mtHSP70 antibodies. Micrographs were acquired on a Zeiss LSM900 confocal microscope with an Airyscan detector. Scale bar = 10 µm. The signal corresponding to each channel along the drawn transect is displayed for each condition in **E**. Representative experiment of two independent biological replicates using two different *MPV17* WT and KO cell lines.

**Expanded View Figure 2.**
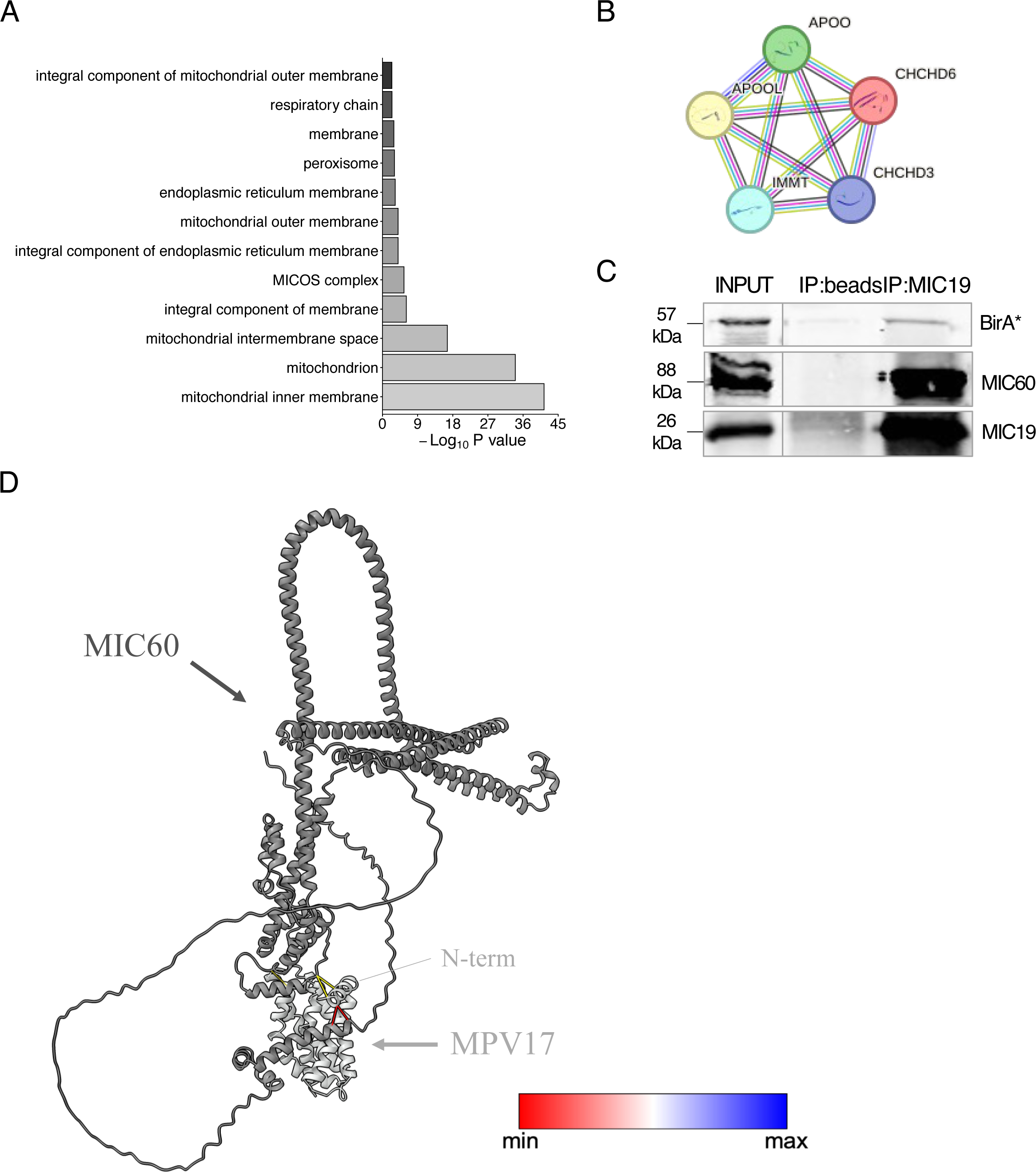
Confirmation of MPV17 interaction with the MICOS and 3D modeling. (**A**) GO enrichment analysis done on the ≥ 2-fold-enriched proteins in the MPV17-BirA* BioID compared to preOTC-BirA* condition, using the DAVID platform and showing the top enriched CC. (**B**) Illustration of the local network cluster corresponding to the “mitochondrial crista junction”, highlighted with the arrow in Figure 3B, and depicting the proteins found in the BioID data associated with this network (IMMT/MIC60, CHCHD3/MIC19, CHCHD6/MIC25, APOO/MIC26 and APOOL/MIC27). (**C**) HEK293T cells were seeded, transfected 24 h later with the MPV17-BirA* construct and harvested 48 h after transfection for protein extraction. 800 µg of proteins were loaded on magnetic beads coupled or not with 5 µg of MIC19 antibody. Proteins were immunoprecipitated for 16 h on a wheel at 4 °C and proteins were resolved by SDS-PAGE. 5% of the lysate (40 µg) was used as input. BirA*, MIC60 and MIC19 were revealed by Western blot analysis (n = 1). (**D**) Representation of the predicted dimerization model of MPV17 (light grey) with MIC60 (dark grey), using ChimeraX software (Meng et al., 2023) and based on the prediction produced with the Alphafold multimer prediction tool. The colored links depict the predicted interaction between amino acids with a confidence level ranging from low (red) to high (blue), showing an interaction of MIC60 with the N-terminal part of MPV17.

**Expanded View Figure 3.**
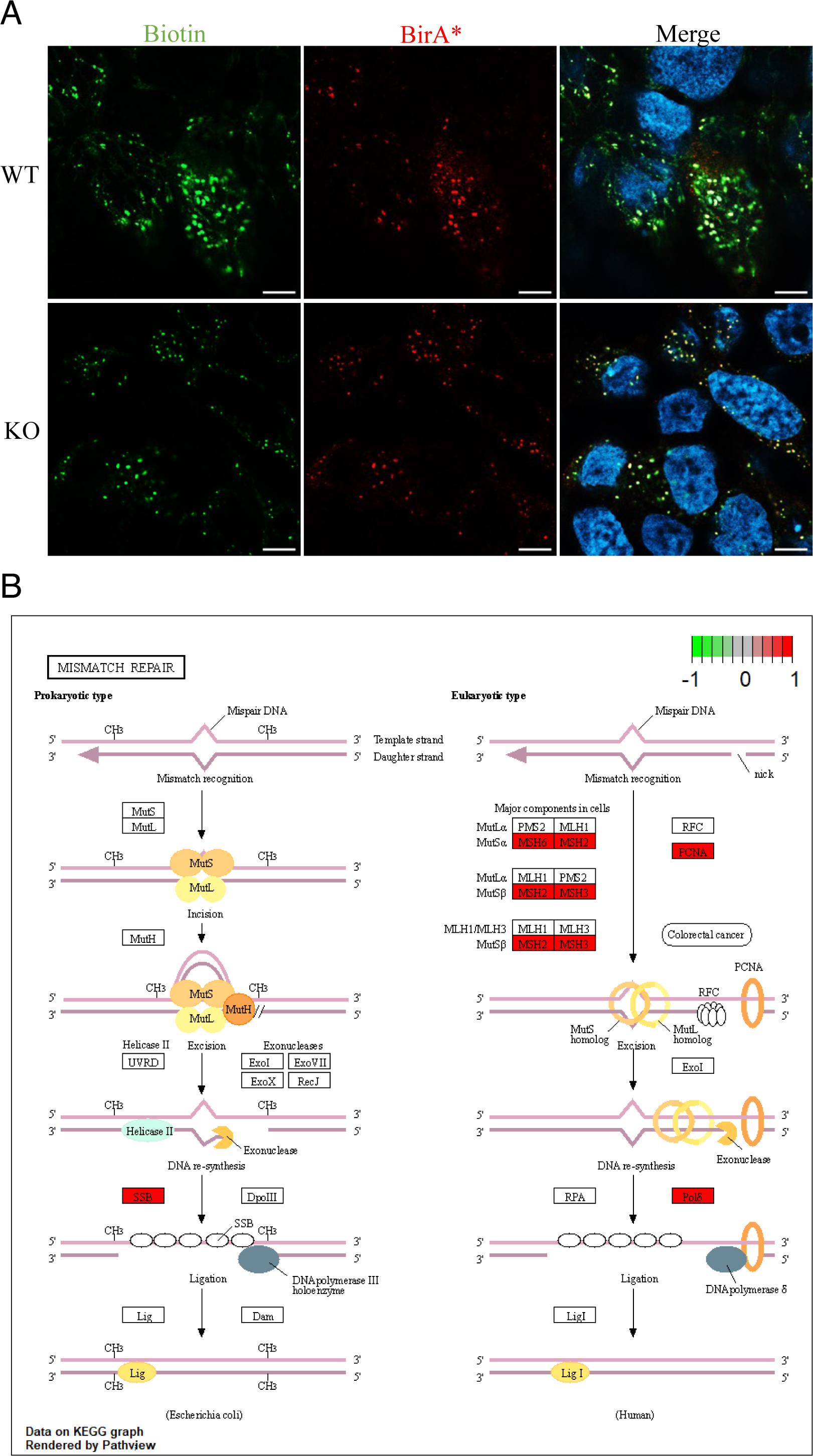
Functional validation of TWNK-BirA* BioID showing enrichment of mismatch repair in KO cells. (**A**) HEK293T*^MPV17+/+^*and HEK293T*^MPV17-/-^* cells were seeded and transfected 24 h later with the TWNK-BirA* construct. The cells were incubated 24 h after with 50 µM of biotin for 6 h before being fixed in 4 % PFA and labelled with anti-BirA* antibody and streptavidin-alexa488 probe to reveal biotinylated proteins. Nuclei were stained using DAPI. Micrographs were acquired on a Zeiss LSM900 confocal microscope with an Airyscan detector. Scale bar = 5 µm. Representative experiment of three independent biological replicates. (**B**) A GSEA was performed, using the KEGG database. The KEGG pathway “Mismatch repair was computed using the “pathview” package (R software) in accordance to the enrichment values (log_2_(fold-change)) of the proteins found in the TWNK-BirA* BioID dataset. The proteins with a green labelling are found negatively enriched whereas the proteins with a red labelling are found positively enriched in the TWNK-BirA* KO condition compared to the TWNK-BirA* WT condition.

## REFERENCES

Achanta, G., Sasaki, R., Feng, L., Carew, J. S., Lu, W., Pelicano, H., Keating, M. J., & Huang, P. (2005). Novel role of p53 in maintaining mitochondrial genetic stability through interaction with DNA Pol γ. EMBO Journal, 24(19), 3482–3492. 10.1038/sj.emboj.7600819

Allen, G. F. G., Toth, R., James, J., & Ganley, I. G. (2013). Loss of iron triggers PINK1/Parkin-independent mitophagy. EMBO Reports, 14(12), 1127–1135. 10.1038/EMBOR.2013.168

Antonenkov, V. D., Isomursu, A., Mennerich, D., Vapola, M. H., Weiher, H., Kietzmann, T., & Kalervo Hiltunen, J. (2015). The Human Mitochondrial DNA Depletion Syndrome Gene MPV17 Encodes a Non-selective Channel That Modulates Membrane Potential *. The Journal of Biological Chemistry, 290. 10.1074/jbc.M114.608083

Ashkenazy, H., Abadi, S., Martz, E., Chay, O., Mayrose, I., Pupko, T., & Ben-Tal, N. (2016). ConSurf 2016: an improved methodology to estimate and visualize evolutionary conservation in macromolecules. Nucleic Acids Research, 44(W1), W344–50. 10.1093/nar/gkw408

Bannwarth, S., Figueroa, A., Fragaki, K., Destroismaisons, L., Lacas-Gervais, S., Lespinasse, F., Vandenbos, F., Pradelli, L. A., Ricci, J.-E., Rötig, A., Michiels, J.-F., Vande Velde, C., & Paquis-Flucklinger, V. (2012). The human MSH5 (MutSHomolog 5) protein localizes to mitochondria and protects the mitochondrial genome from oxidative damage. Mitochondrion, 12(6), 654–665. 10.1016/j.mito.2012.07.111

Bergant, V., Schnepf, D., de Andrade Krätzig, N., Hubel, P., Urban, C., Engleitner, T., Dijkman, R., Ryffel, B., Steiger, K., Knolle, P. A., Kochs, G., Rad, R., Staeheli, P., & Pichlmair, A. (2023). mRNA 3’UTR lengthening by alternative polyadenylation attenuates inflammatory responses and correlates with virulence of Influenza A virus. Nature Communications, 14(1), 4906. 10.1038/s41467-023-40469-6

Bernardi, P., Gerle, C., Halestrap, A. P., Jonas, E. A., Karch, J., & Mnatsakanyan, N. (2023). Identity, structure, and function of the mitochondrial permeability transition pore□: controversies, consensus, recent advances, and future directions. CDDpress, 30(June), 1869–1885. 10.1038/s41418-023-01187-0

Bian, W., Pu, S., Xie, S., Wang, C., Deng, S., Strauss, P. R., & Pei, D. (2021). Loss of mpv17 affected early embryonic development via mitochondria dysfunction in zebra fi sh. Cell Death Discovery, (August), 1–10. 10.1038/s41420-021-00630-w

Blakely, E. L., Butterworth, A., Hadden, R. D. M., Bodi, I., He, L., McFarland, R., & Taylor, R. W. (2012). MPV17 mutation causes neuropathy and leukoencephalopathy with multiple mtDNA deletions in muscle. Neuromuscular Disorders, 22(7), 587–591. 10.1016/J.NMD.2012.03.006

Blighe, K., Rana, S., Turkes, E., Ostenforf, B., Grioni, A., & Lewis, M. (2023). Publication-ready volcano plots with enhanced colouring and labelling. 10.18129/B9.bioc.EnhancedVolcano

Bottani, E., Giordano, C., Civiletto, G., Di Meo, I., Auricchio, A., Ciusani, E., Marchet, S., Lamperti, C., D’Amati, G., Viscomi, C., & Zeviani, M. (2014). AAV-mediated Liver-specific MPV17 Expression Restores mtDNA Levels and Prevents Diet-induced Liver Failure. Molecular Therapy, 22(1), 10–17. 10.1038/MT.2013.230

Brookes, P. S., Yoon, Y., Robotham, J. L., Anders, M. W., & Sheu, S. S. (2004). Calcium, ATP, and ROS: A mitochondrial love-hate triangle. American Journal of Physiology - Cell Physiology, 287(4 56-4). 10.1152/ajpcell.00139.2004

Canonne, M., Wanet, A., Nguyen, T. T. A., Khelfi, A., Ayama-Canden, S., Van Steenbrugge, M., Fattaccioli, A., Sokal, E., Najimi, M., Arnould, T., & Renard, P. (2020). MPV17 does not control cancer cell proliferation. PLOS ONE, 15(3), e0229834. 10.1371/journal.pone.0229834

Choi, Y., Hong, Y. Bin, Jung, S., Lee, J. H., Kim, Y. J., Park, H. J., Lee, J., Koo, H., Lee, J., Jwa, D. H., Jung, N., Woo, S., Kim, S., & Chung, K. W. (2015). A novel homozygous MPV17 mutation in two families with axonal sensorimotor polyneuropathy. BMC Neurology, 1–10. 10.1186/s12883-015-0430-1

Concordet, J. P., & Haeussler, M. (2018). CRISPOR: intuitive guide selection for CRISPR/Cas9 genome editing experiments and screens. Nucleic Acids Research, 46(W1), W242–W245. 10.1093/NAR/GKY354

Corra, S., Viscomi, C., & Costa, R. (2023). Drosophila Mpv17 forms an ion channel and regulates energy metabolism. IScience, 26(107955). 10.1016/j.isci.2023.107955

Creed, T. M., Baldeosingh, R., Eberly, C. L., Schlee, C. S., Kim, M., Cutler, J. A., Pandey, A., Civin, C. I., Fossett, N. G., & Kingsbury, T. J. (2020). The PAX-SIX-EYA-DACH network modulates GATA-FOG function in fly hematopoiesis and human erythropoiesis. Development (Cambridge, England), 147(1). 10.1242/dev.177022

Cullen, P. J., & Steinberg, F. (2018). To degrade or not to degrade: mechanisms and significance of endocytic recycling. Nature Reviews Molecular Cell Biology 2018 19:11, 19(11), 679–696. 10.1038/s41580-018-0053-7

Dalla Rosa, I., Cámara, Y., Durigon, R., Moss, C. F., Vidoni, S., Akman, G., Hunt, L., Johnson, M. A., Grocott, S., Wang, L., Thorburn, D. R., Hirano, M., Poulton, J., Taylor, R. W., Elgar, G., Martí, R., Voshol, P., … Spinazzola, A. (2016). MPV17 Loss Causes Deoxynucleotide Insufficiency and Slow DNA Replication in Mitochondria. PLOS Genetics, 12(1), e1005779. 10.1371/journal.pgen.1005779

Dallabona, C., Marsano, R. M., Arzuffi, P., Ghezzi, D., Mancini, P., Zeviani, M., Ferrero, I., & Donnini, C. (2010). Sym1, the yeast ortholog of the MPV17 human disease protein, is a stress-induced bioenergetic and morphogenetic mitochondrial modulator. Human Molecular Genetics, 19(6), 1098–1107. 10.1093/hmg/ddp581

Dennis, G., Sherman, B. T., Hosack, D. A., Yang, J., Gao, W., Lane, H. C., & Lempicki, R. A. (2003). DAVID: Database for Annotation, Visualization, and Integrated Discovery. Genome Biology, 4(5), 1–11. 10.1186/GB-2003-4-9-R60/TABLES/3

Denton, R. M., Richards, D. A., & Chin, J. G. (1978). Calcium ions and the regulation of NAD+-linked isocitrate dehydrogenase from the mitochondria of rat heart and other tissues. The Biochemical Journal, 176(3), 899–906. 10.1042/bj1760899

El-Hattab, A., Li, F., Schmitt, E., Zhang, S., Craigen, W., & Wong, L. (2010). MPV17-associated hepatocerebral mitochondrial DNA depletion syndrome: new patients and novel mutations. Molecular Genetics and Metabolism, 99(3), 300–308. 10.1016/J.YMGME.2009.10.003

El-Hattab, A. W., Craigen, W. J., & Scaglia, F. (2017). Mitochondrial DNA maintenance defects. Biochimica et Biophysica Acta (BBA) - Molecular Basis of Disease, 1863(6), 1539–1555. 10.1016/J.BBADIS.2017.02.017

El-Hattab, A. W., Wang, J., Dai, H., Almannai, M., Staufner, C., Alfadhel, M., Gambello, M. J., Prasun, P., Raza, S., Lyons, H. J., Afqi, M., Saleh, M. A. M., Faqeih, E. A., Alzaidan, H. I., Alshenqiti, A., Flore, L. A., Hertecant, J., … Wong, L. J. C. (2018). MPV17-related mitochondrial DNA maintenance defect: New cases and review of clinical, biochemical, and molecular aspects. Human Mutation, 39(4), 461–470. 10.1002/humu.23387

Elfmann, C., & Stülke, J. (2023). PAE viewer: A webserver for the interactive visualization of the predicted aligned error for multimer structure predictions and crosslinks. Nucleic Acids Research, 51(W1), W404–W410. 10.1093/nar/gkad350

Evans, R., O’Neill, M., Pritzel, A., Antropova, N., Senior, A. W., Green, T., Žídek, A., Bates, R., Blackwell, S., Yim, J., Ronneberger, O., Bodenstein, S., Zielinski, M., Bridgland, A., Potapenko, A., Cowie, A., Tunyasuvunakool, K., … Hassabis, D. (2022). AlphaFold-Multimer Protein complex prediction with. BioRxiv. 10.1101/2021.10.04.463034

Fernandez-Sanz, C., De la Fuente, S., & Sheu, S. S. (2019). Mitochondrial Ca2+ concentrations in live cells: quantification methods and discrepancies. FEBS Letters, 593(13), 1528–1541. 10.1002/1873-3468.13427

Fontana, G. A., & Gahlon, H. L. (2020, November 18). Mechanisms of replication and repair in mitochondrial DNA deletion formation. Nucleic Acids Research. Oxford University Press. 10.1093/nar/gkaa804

Friese, C., Yang, J., Mendelsohn-Victor, K., & Mccullagh, M. (2019). The “fast” and the “slow” modes of mitochondrial DNA degradation. Physiology & Behavior, 46(2), 248–256. 10.3109/19401736.2014.905829.The

Fu, Y., Tigano, M., & Sfeir, A. (2020). Safeguarding mitochondrial genomes in higher eukaryotes. Nature Structural & Molecular Biology 2020 27:8, 27(8), 687–695. 10.1038/s41594-020-0474-9

Gilberti, M., Baruffini, E., Donnini, C., & Dallabona, C. (2018). Pathological alleles of MPV17 modeled in the yeast Saccharomyces cerevisiae orthologous gene SYM1 reveal their inability to take part in a high molecular weight complex. PLOS ONE, 13(10), e0205014. 10.1371/JOURNAL.PONE.0205014

Giorgio, V., von Stockum, S., Antoniel, M., Fabbro, A., Fogolari, F., Forte, M., Glick, G. D., Petronilli, V., Zoratti, M., Szabó, I., Lippe, G., & Bernardi, P. (2013). Dimers of mitochondrial ATP synthase form the permeability transition pore. Proceedings of the National Academy of Sciences of the United States of America, 110(15), 5887–5892. 10.1073/pnas.1217823110

Han, S., Udeshi, N. D., Deerinck, T. J., Svinkina, T., Ellisman, M. H., Carr, S. A., & Ting, A. Y. (2017). Proximity Biotinylation as a Method for Mapping Proteins Associated with mtDNA in Living Cells. Cell Chemical Biology, 24(3), 404–414. 10.1016/j.chembiol.2017.02.002

Hurst, S., Hoek, J., & Sheu, S. S. (2017). Mitochondrial Ca2+ and regulation of the permeability transition pore. Journal of Bioenergetics and Biomembranes, 49(1), 27–47. 10.1007/s10863-016-9672-x

Jacinto, S., Guerreiro, P., de Oliveira, R. M., Cunha-Oliveira, T., Santos, M. J., Grazina, M., Rego, A. C., & Outeiro, T. F. (2021). MPV17 Mutations Are Associated With a Quiescent Energetic Metabolic Profile. Frontiers in Cellular Neuroscience, 0, 47. 10.3389/FNCEL.2021.641264

Jia, J., Xia, J., Liu, W., Tao, F., & Xiao, J. (2023). Cinnamtannin B-1 Inhibits the Progression of Osteosarcoma by Regulating the miR-1281/PPIF Axis. Biological & Pharmaceutical Bulletin, 46(1), 67–73. 10.1248/bpb.b22-00600

Jumper, J., Evans, R., Pritzel, A., Green, T., Figurnov, M., Ronneberger, O., Tunyasuvunakool, K., Bates, R., Žídek, A., Potapenko, A., Bridgland, A., Meyer, C., Kohl, S. A. A., Ballard, A. J., Cowie, A., Romera-Paredes, B., Nikolov, S., … Hassabis, D. (2021). Highly accurate protein structure prediction with AlphaFold. Nature, 596(7873), 583–589. 10.1038/s41586-021-03819-2

Kalifa, L., Quintana, D. F., Schiraldi, L. K., Phadnis, N., Coles, G. L., Sia, R. A., & Sia, E. A. (2012). Mitochondrial genome maintenance: Roles for nuclear nonhomologous end-joining proteins in Saccharomyces cerevisiae. Genetics, 190(3), 951–964. 10.1534/genetics.111.138214

Karch, J., Bround, M. J., Khalil, H., Sargent, M. A., Latchman, N., Terada, N., Peixoto, P. M., & Molkentin, J. D. (2019). Inhibition of mitochondrial permeability transition by deletion of the ANT family and CypD. Science Advances, 5(8), 1–7. 10.1126/sciadv.aaw4597

Killilea, D. W., & Killilea, A. N. (2022). Mineral requirements for mitochondrial function: A connection to redox balance and cellular differentiation. Free Radical Biology and Medicine, 182(October 2021), 182–191. 10.1016/j.freeradbiomed.2022.02.022

Kim, J., Cantor, A. B., Orkin, S. H., & Wang, J. (2009). Use of in vivo biotinylation to study protein–protein and protein–DNA interactions in mouse embryonic stem cells. Nature Protocols, 4(4), 506–517. 10.1038/nprot.2009.23

Kondadi, A. K., Anand, R., Hänsch, S., Urbach, J., Zobel, T., Wolf, D. M., Segawa, M., Liesa, M., Shirihai, O. S., Weidtkamp-Peters, S., & Reichert, A. S. (2020). Cristae undergo continuous cycles of membrane remodelling in a MICOS-dependent manner. EMBO Reports, 21(3), e49776. 10.15252/embr.201949776

Kowaltowski, A. J., Castilho, R. F., & Vercesi, A. E. (1996). Opening of the mitochondrial permeability transition pore by uncoupling or inorganic phosphate in the presence of Ca2+ is dependent on mitochondrial-generated reactive oxygen species. FEBS Letters, 378(2), 150–152. 10.1016/0014-5793(95)01449-7

Kozjak-Pavlovic, V. (2017, January 1). The MICOS complex of human mitochondria. Cell and Tissue Research. Springer Verlag. 10.1007/s00441-016-2433-7

Kwon, S.-K., Sando, R. 3rd, Lewis, T. L., Hirabayashi, Y., Maximov, A., & Polleux, F. (2016). LKB1 Regulates Mitochondria-Dependent Presynaptic Calcium Clearance and Neurotransmitter Release Properties at Excitatory Synapses along Cortical Axons. PLoS Biology, 14(7), e1002516. 10.1371/journal.pbio.1002516

Le Sage, V., Cinti, A., & Mouland, A. J. (2016). Proximity□Dependent Biotinylation for Identification of Interacting Proteins. Current Protocols in Cell Biology, 73(1), 17.19.1-17.19.12. 10.1002/cpcb.11

Lin, H., He, L., & Ma, B. (2013). A combinatorial approach to the peptide feature matching problem for label-free quantification. Bioinformatics (Oxford, England), 29(14), 1768–1775. 10.1093/BIOINFORMATICS/BTT274

Liu, X., Salokas, K., Tamene, F., Jiu, Y., Weldatsadik, R. G., Öhman, T., & Varjosalo, M. (2018). An AP-MS-and BioID-compatible MAC-tag enables comprehensive mapping of protein interactions and subcellular localizations. Nature Communications, 1188(9). 10.1038/s41467-018-03523-2

Luna-Sanchez, M., Benincá, C., Cerutti, R., Brea-Calvo, G., Yeates, A., Scorrano, L., Zeviani, M., & Viscomi, C. (2020). Opa1 Overexpression Protects from Early-Onset Mpv17−/−-Related Mouse Kidney Disease. Molecular Therapy, 28(8), 1918–1930. 10.1016/j.ymthe.2020.06.010

Luo, W., & Brouwer, C. (2013). Pathview: an R/Bioconductor package for pathway-based data integration and visualization. Bioinformatics, 29(14), 1830–1831. doi:10.1093/bioinformatics/btt285

Madungwe, N. B., Feng, Y., Aliagan, A. I., Tombo, N., Kaya, F., & Bopassa, J. C. (2020). Inner mitochondrial membrane protein MPV17 mutant mice display increased myocardial injury after ischemia/reperfusion. American Journal of Translational Research, 12(7), 3412. Retrieved from /pmc/articles/PMC7407695/

Masood, T. Bin, Sandhya, S., Chandra, N., & Natarajan, V. (2015). CHEXVIS: A tool for molecular channel extraction and visualization. BMC Bioinformatics, 16(1), 1–19. 10.1186/s12859-015-0545-9

Mattingly, R. R., & Macara, I. G. (1996). Phosphorylation-dependent activation of the Ras-GRF/CDC25Mm exchange factor by muscarinic receptors and G-protein beta gamma subunits. Nature, 382(6588), 268–272. 10.1038/382268a0

McCormack, J. G., & Denton, R. M. (1979). The effects of calcium ions and adenine nucleotides on the activity of pig heart 2-oxoglutarate dehydrogenase complex. The Biochemical Journal, 180(3), 533–544. 10.1042/bj1800533

Meier, F., Brunner, A. D., Koch, S., Koch, H., Lubeck, M., Krause, M., Goedecke, N., Decker, J., Kosinski, T., Park, M. A., Bache, N., Hoerning, O., Cox, J., Räther, O., & Mann, M. (2018). Online Parallel Accumulation-Serial Fragmentation (PASEF) with a Novel Trapped Ion Mobility Mass Spectrometer. Molecular & Cellular ProteomicslJ: MCP, 17(12), 2534–2545. 10.1074/MCP.TIR118.000900

Meng, E. C., Goddard, T. D., Pettersen, E. F., Couch, G. S., Pearson, Z. J., Morris, J. H., & Ferrin, T. E. (2023). UCSF ChimeraX: Tools for structure building and analysis. Protein Science, 32(11), 1–13. 10.1002/pro.4792

Meurant, S., Mauclet, L., Dieu, M., Arnould, T., Eyckerman, S., & Renard, P. (2023). Endogenous TOM20 Proximity Labelling: A Swiss-Knife for the Study of Mitochondrial Proteins in Human Cells. International Journal of Molecular Sciences, 24(11). 10.3390/ijms24119604

Mnatsakanyan, N., Llaguno, M. C., Yang, Y., Yan, Y., Weber, J., Sigworth, F. J., & Jonas, E. A. (2019). A mitochondrial megachannel resides in monomeric F1FO ATP synthase. Nature Communications, 10(1). 10.1038/s41467-019-13766-2

Morgan, A. J., & Jacob, R. (1994). Ionomycin enhances Ca2+ influx by stimulating store-regulated cation entry and not by a direct action at the plasma membrane. The Biochemical Journal, 300 (Pt 3(Pt 3), 665–672. 10.1042/bj3000665

Mukherjee, S., Das, S., Bedi, M., Vadupu, L., Ball, W. B., & Ghosh, A. (2023). Methylglyoxal-mediated Gpd1 activation restores the mitochondrial defects in a yeast model of mitochondrial DNA depletion syndrome. Biochimica et Biophysica Acta - General Subjects, 1867(5), 130328. 10.1016/j.bbagen.2023.130328

Naumova, N., & Šachl, R. (2020). Regulation of cell death by mitochondrial transport systems of calcium and BCL□2 proteins. Membranes, 10(10), 1–33. 10.3390/membranes10100299

Nicolas, E., Simion, P., Guérineau, M., Terwagne, M., Colinet, M., Virgo, J., Lingurski, M., Boutsen, A., Dieu, M., Hallet, B., & Van Doninck, K. (2023). Horizontal acquisition of a DNA ligase improves DNA damage tolerance in eukaryotes. Nature Communications, 14(1). 10.1038/s41467-023-43075-8

Nissanka, N., Bacman, S. R., Plastini, M. J., & Moraes, C. T. (2018). The mitochondrial DNA polymerase gamma degrades linear DNA fragments precluding the formation of deletions. Nature Communications, 9(1), 1–9. 10.1038/s41467-018-04895-1

Pankiv, S., Clausen, T. H., Lamark, T., Brech, A., Bruun, J.-A., Outzen, H., Øvervatn, A., Bjørkøy, G., & Johansen, T. (2007). p62/SQSTM1 Binds Directly to Atg8/LC3 to Facilitate Degradation of Ubiquitinated Protein Aggregates by Autophagy *. Journal of Biological Chemistry, 282(33), 24131–24145. 10.1074/JBC.M702824200

Papeta, N., Zheng, Z., Schon, E. A., Brosel, S., Altintas, M. M., Nasr, S. H., Reiser, J., D’Agati, V. D., & Gharavi, A. G. (2010). Prkdc participates in mitochondrial genome maintenance and prevents adriamycin-induced nephropathy in mice. Journal of Clinical Investigation, 120(11), 4055–4064. 10.1172/JCI43721

Peeva, V., Blei, D., Trombly, G., Corsi, S., Szukszto, M. J., Rebelo-Guiomar, P., Gammage, P. A., Kudin, A. P., Becker, C., Altmüller, J., Minczuk, M., Zsurka, G., & Kunz, W. S. (2018). Linear mitochondrial DNA is rapidly degraded by components of the replication machinery. Nature Communications, 9(1), 1–11. 10.1038/s41467-018-04131-w

Peng, T.-I., & Greenamyre, J. T. (1998). Privileged Access to Mitochondria of Calcium Influx through N-Methyl-d-Aspartate Receptors. Molecular Pharmacology, 53(6), 974 LP – 980. Retrieved from http://molpharm.aspetjournals.org/content/53/6/974.abstract

Perez-Riverol, Y., Bai, J., Bandla, C., García-Seisdedos, D., Hewapathirana, S., Kamatchinathan, S., Kundu, D. J., Prakash, A., Frericks-Zipper, A., Eisenacher, M., Walzer, M., Wang, S., Brazma, A., & Vizcaíno, J. A. (2022). The PRIDE database resources in 2022: a hub for mass spectrometry-based proteomics evidences. Nucleic Acids Research, 50(D1), D543–D552. 10.1093/NAR/GKAB1038

Picca, A., Guerra, F., Calvani, R., Coelho-Junior, H. J., Bossola, M., Landi, F., Bernabei, R., Bucci, C., & Marzetti, E. (2020). Generation and Release of Mitochondrial-Derived Vesicles in Health, Aging and Disease. Journal of Clinical Medicine, 9(5). 10.3390/JCM9051440

Quiles, J. M., & Gustafsson, Å. B. (2020). Mitochondrial Quality Control and Cellular Proteostasis: Two Sides of the Same Coin. Frontiers in Physiology, 11, 515. 10.3389/FPHYS.2020.00515/BIBTEX

Rashid, S., Freitas, M. O., Cucchi, D., Bridge, G., Yao, Z., Gay, L., Williams, M., Wang, J., Suraweera, N., Silver, A., McDonald, S. A. C., Chelala, C., Szabadkai, G., & Martin, S. A. (2019). MLH1 deficiency leads to deregulated mitochondrial metabolism. Cell Death and Disease, 10(11). 10.1038/s41419-019-2018-y

Rong, Z., Tu, P., Xu, P., Sun, Y., Yu, F., Tu, N., Guo, L., & Yang, Y. (2021). The Mitochondrial Response to DNA Damage. Frontiers in Cell and Developmental Biology, 9(May), 1–10. 10.3389/fcell.2021.669379

Roux, K. J., Kim, D. I., Raida, M., & Burke, B. (2012). A promiscuous biotin ligase fusion protein identifies proximal and interacting proteins in mammalian cells. Journal of Cell Biology, 196(6), 801–810. 10.1083/jcb.201112098

Sage, J. M., Gildemeister, O. S., & Knight, K. L. (2010). Discovery of a novel function for human Rad51: Maintenance of the mitochondrial genome. Journal of Biological Chemistry, 285(25), 18984–18990. 10.1074/jbc.M109.099846

Sen, A., Kallabis, S., Gaedke, F., Jüngst, C., Boix, J., Nüchel, J., Maliphol, K., Hofmann, J., Schauss, A. C., Krüger, M., Wiesner, R. J., & Pla-Martín, D. (2022). Mitochondrial membrane proteins and VPS35 orchestrate selective removal of mtDNA. Nature Communications 2022 13:1, 13(1), 1–20. 10.1038/s41467-022-34205-9

Shu, L., Hu, C., Xu, M., Yu, J., He, H., Lin, J., Sha, H., Lu, B., Engelender, S., Guan, M., & Song, Z. (2021). ATAD3B is a mitophagy receptor mediating clearance of oxidative stress□induced damaged mitochondrial DNA. The EMBO Journal, 40(8). 10.15252/EMBJ.2020106283

Sohn, J. H., Mutlu, B., Latorre-Muro, P., Liang, J., Bennett, C. F., Sharabi, K., Kantorovich, N., Jedrychowski, M., Gygi, S. P., Banks, A. S., & Puigserver, P. (2023). Liver mitochondrial cristae organizing protein MIC19 promotes energy expenditure and pedestrian locomotion by altering nucleotide metabolism. Cell Metabolism, 35(8), 1356–1372.e5. 10.1016/j.cmet.2023.06.015

Sperl, L. E., & Hagn, F. (2021). NMR Structural and Biophysical Analysis of the Disease-Linked Inner Mitochondrial Membrane Protein MPV17. Journal of Molecular Biology, 433(15), 167098. 10.1016/J.JMB.2021.167098

Spinazzola, A., Viscomi, C., Fernandez-Vizarra, E., Carrara, F., D’Adamo, P., Calvo, S., Marsano, R. M., Donnini, C., Weiher, H., Strisciuglio, P., Parini, R., Sarzi, E., Chan, A., DiMauro, S., Rötig, A., Gasparini, P., Ferrero, I., … Zeviani, M. (2006). MPV17 encodes an inner mitochondrial membrane protein and is mutated in infantile hepatic mitochondrial DNA depletion. Nature Genetics, 38(5), 570–576. 10.1038/ng1765

Stewart, J. D., Schoeler, S., Sitarz, K. S., Horvath, R., Hallmann, K., Pyle, A., Yu-Wai-Man, P., Taylor, R. W., Samuels, D. C., Kunz, W. S., & Chinnery, P. F. (2011). POLG mutations cause decreased mitochondrial DNA repopulation rates following induced depletion in human fibroblasts. Biochimica et Biophysica Acta - Molecular Basis of Disease, 1812(3), 321–325. 10.1016/j.bbadis.2010.11.012

Sugiura, A., McLelland, G., Fon, E. A., & McBride, H. M. (2014). A new pathway for mitochondrial quality control: mitochondrial□derived vesicles. The EMBO Journal, 33(19), 2142–2156. 10.15252/EMBJ.201488104

Suomalainen, A., & Isohanni, P. (2010). Mitochondrial DNA depletion syndromes – Many genes, common mechanisms. Neuromuscular Disorders, 20(7), 429–437. 10.1016/J.NMD.2010.03.017

Suzek, B. E., Wang, Y., Huang, H., McGarvey, P. B., & Wu, C. H. (2015). UniRef clusters: a comprehensive and scalable alternative for improving sequence similarity searches. Bioinformatics (Oxford, England), 31(6), 926–932. 10.1093/bioinformatics/btu739

Tanveer, A., Virji, S., Andreeva, L., Totty, N. F., Hsuan, J. J., Ward, J. M., & Crompton, M. (1996). Involvement of cyclophilin D in the activation of a mitochondrial pore by Ca2+ and oxidant stress. European Journal of Biochemistry, 238(1), 166–172. 10.1111/j.1432-1033.1996.0166q.x

Tanwar, J., Singh, J. B., & Motiani, R. K. (2021). Molecular machinery regulating mitochondrial calcium levels: The nuts and bolts of mitochondrial calcium dynamics. Mitochondrion, 57, 9–22. 10.1016/J.MITO.2020.12.001

Todkar, K., Chikhi, L., Desjardins, V., El-Mortada, F., Pépin, G., & Germain, M. (2021). Selective packaging of mitochondrial proteins into extracellular vesicles prevents the release of mitochondrial DAMPs. Nature Communications 2021 12:1, 12(1), 1–12. 10.1038/s41467-021-21984-w

Urbani, A., Giorgio, V., Carrer, A., Franchin, C., Arrigoni, G., Jiko, C., Abe, K., Maeda, S., Shinzawa-Itoh, K., Bogers, J. F. M., McMillan, D. G. G., Gerle, C., Szabò, I., & Bernardi, P. (2019). Purified F-ATP synthase forms a Ca(2+)-dependent high-conductance channel matching the mitochondrial permeability transition pore. Nature Communications, 10(1), 4341. 10.1038/s41467-019-12331-1

von Mering, C., Huynen, M., Jaeggi, D., Schmidt, S., Bork, P., & Snel, B. (2003). STRING: a database of predicted functional associations between proteins. Nucleic Acids Research, 31(1), 258–261. Retrieved from http://www.ncbi.nlm.nih.gov/pubmed/12519996

Wang, K., Huang, C., Jiang, T., Chen, Z., Xue, M., Zhang, Q., Zhang, J., & Dai, J. (2021). RNA-binding protein RBM47 stabilizes IFNAR1 mRNA to potentiate host antiviral activity. EMBO Reports, 22(8), e52205. 10.15252/embr.202052205

Werley, C. A., Boccardo, S., Rigamonti, A., Hansson, E. M., & Cohen, A. E. (2020). Multiplexed Optical Sensors in Arrayed Islands of Cells for multimodal recordings of cellular physiology. Nature Communications, 11(1), 3881. 10.1038/s41467-020-17607-5

Wu, T., Hu, E., Xu, S., Chen, M., Guo, P., Dai, Z., Feng, T., Zhou, L., Tang, W., Zhan, L., Fu, X., Liu, S., Bo, X., & Yu, G. (2021). clusterProfiler 4.0: A universal enrichment tool for interpreting omics data. The Innovation, 2(3). 10.1016/J.XINN.2021.100141

Yu, G. (2021). enrichplot: Visualization of Functional Enrichment Result. doi:10.18129/B9.bioc.enrichplot

Zaman, Q., Khan, M. A., Sahar, K., Rehman, G., Khan, H., Rehman, M., Ahmad, I., Tariq, M., Muthaffar, O. Y., Abdulkareem, A. A., Bibi, F., Naseer, M. I., Faisal, M. S., Wasif, N., & Jelani, M. (2023). Charcot – Marie – Tooth and Spastic Ataxia of Charlevoix – Saguenay Type Diseases. Genes, 14(328), 1–15.

Zhao, L. (2019). Mitochondrial DNA degradation: A quality control measure for mitochondrial genome maintenance and stress response. Enzymes (1st ed., Vol. 45). Elsevier Inc. 10.1016/bs.enz.2019.08.004

Zhu, L. P., Yu, X. D., Ling, S., Brown, R. A., & Kuo, T. H. (2000). Mitochondrial Ca(2+)homeostasis in the regulation of apoptotic and necrotic cell deaths. Cell Calcium, 28(2), 107–117. 10.1054/ceca.2000.0138

